# T4-type phages diversity in wetland soils reveals their ubiquity and their likely host-dependent dynamics

**DOI:** 10.64898/2026.07.13.738189

**Authors:** Rémi Trémouille, Virginie Daburon, Achim Quaiser, Alexis Dufresne, Cécile Monard

## Abstract

Bacteriophages are abundant and diverse in soils, playing a major role in regulating bacterial communities and consequently affecting biogeochemical cycles. Such host-phage interactions may be influenced by fluctuations in soil moisture, as observed in wetlands soils which constitute a key feature of the ongoing climate change. Here, we investigated the spatial and temporal dynamics of both bacteria and T4-type bacteriophage community structures and diversities in soil of a freshwater wetland. Soil was sampled in three sites across a proximal soil transect presenting an increase moisture content at seven dates over an 18 months period with contrasted flooding periods. DNA was extracted and we applied amplicon sequencing of the bacterial 16S rRNA gene and viral g23 gene. Bacterial community composition varied across the proximal soil transect, with *Methylomirabilia* and *Gammaproteobacteria* being significantly enriched in the wettest site and comprising ASVs affiliated to methanotroph and denitrifying bacteria, respectively. We identified a large diversity of T4-type phages, among which a fraction was novel, while others were similar to phages previously sequenced from various biomes. These findings suggest that T4-type phages are capable of successfully colonizing diverse niches in the biosphere, contributing to their ubiquity and diversity. Viral community was however dominated by few vASVs, which were highly represented in one or two of the three studied sites supporting the Bank model. All together our results indicate that T4-type phages have broad host ranges and more likely follow bacterial population dynamics. The present study provides new insights into the role of phages in soil, highlighting their interactions with bacterial hosts involved in carbon and nitrogen cycles, interactions that are likely regulated by fluctuations in soil moisture, as observed in wetlands.

**Highlights:**

- Both bacterial and T4-type phages were structured across proximal sites
- Bacterial 16S rRNA gene copy number was inversely correlated to the soil moisture
- 26 viral ASVs did not cluster with reference sequences
- viral ASVs seem to be primarily controlled by host availability
- Soil bacteria and phage diversities were significantly lower in the wettest site

## 1. Introduction

Bacteriophages (‘Phages’), viruses infecting bacteria, are the most abundant biological entities on earth, playing key roles in shaping microbial communities and consequently influencing biogeochemical cycles (Hendrix *et al*., 1999; Weinbauer, 2004; Suttle, 2005). Phages and their hosts have intricate interplays combining antagonistic and mutualistic dynamics. Through host lysis, phages exert a top-down control on bacterial populations and promote microbial biomass turnover by releasing nutrients trapped in microbial biomass (*i.e.* the viral shunt; Wilhelm and Suttle, 1999). Such changes in resources availability induce indirect bottom-up controls of bacterial communities (Chow *et al*., 2014).

In soil, bacteria are highly abundant and diverse, constituting a significant reservoir of phage hosts. Since soil bacteria are key actors of soil functioning, their interplay with phages is of importance in soil ecology, driving nutrient cycling and modulating soil health (Kuzyakov and Mason-Jones, 2018; Starr *et al*., 2019; Braga *et al*., 2020; Wang *et al*., 2024; Zhou *et al*., 2024). Soil viral diversity is influenced by several abiotic and biotic soil properties such as texture, moisture, pH as well as the presence of plants (Williamson *et al*., 2017; Emerson *et al*., 2018; Bi *et al*., 2021; Ma *et al*., 2024). Phage-host interactions rely on environmental conditions and soil properties, which affect bacterial activity and regulate encounter rates. These interactions occur at the scale of the soil microbial habitats, which are composed of heterogeneous aggregates and pore spaces (Erktan *et al*., 2020). In favorable conditions, these microniches constitute hot spots and hot moments for phage predation, affecting in turn bacterial diversity (Pratama and van Elsas, 2018).

Within the large diversity of known phages, T4-type phages are tailed dsDNA phages, important members of the *Caudoviricetes* class from the *Straboviridae* family (formerly *Myoviridae*) and widely distributed in diverse natural environments (Desplats and Krisch, 2003; Filée *et al*., 2005; López-Bueno *et al*., 2009; Fujihara *et al*., 2010; Li *et al*., 2019a). Their study often relies on the conserved g23 viral marker gene encoding the major capsid protein (Tetart *et al*., 2001; Desplats and Krisch, 2003; Petrov *et al*., 2010). Since the design of specific primers to study marine T4-type phages (Filée *et al*., 2005), the g23 viral marker has been used on diverse environments such as freshwater (Dar *et al*., 2023), glacier (Bellas and Anesio, 2013), rice paddy soil and water (Fujii *et al*., 2008; Wang *et al*., 2009b; Cahyani *et al*., 2009; Liu *et al*., 2012) or unsaturated soil (Dar *et al*., 2023). Initially these studies were based on clones sequencing (Filée *et al*., 2005), limiting the amount of T4-type phage sequences, but recently, Dar et al. (2023) and Cai et al. (2023) successfully applied high throughput sequencing (HTS) approaches to generate g23 operational taxonomic units (OTUs).

The use of g23 viral marker to study T4-type phages revealed that these viruses are ubiquitous and diverse, implying a flexibility to adapt to new environments and new hosts that led them to successfully expand their niche in the biosphere (Krisch and Comeau, 2008). T4-type phages infect a broad range of bacteria including various genera of *Pseudomonadota* and *Cyanobacteria* (Nakayama *et al*., 2009; Petrov *et al*., 2010; Sullivan *et al*., 2010; Roux *et al*., 2015). Such broad host range should imply that these viruses largely regulate bacterial dynamics in soil through top-down control, playing a key role in soil functioning and biogeochemical cycles.

Host-phage interactions may be influenced by fluctuations in soil moisture, as observed in wetlands soils which constitute a key feature of the ongoing climate change. While their anoxic conditions participate in slowing down the soil organic matter (SOM) decomposition and lead to its accumulation, in return, wetland soil systems provide optimum conditions for methanogenesis (Lyu *et al*., 2018) and denitrification (Hu *et al*., 2015). Here, we investigated the spatial and temporal dynamics of both bacteria and T4-type bacteriophage community structures and diversities in soil of a freshwater wetland. We hypothesized that both soil viral and bacterial diversities i) differ across a proximal soil transect, even within close habitats, and ii) respond to soil moisture dictated by seasonal changes in wetland water table level. We also aimed to identify viral amplicon sequence variants (ASVs) which co-occured more frequently with bacterial ASVs suggesting host-phage relationships. Soil was sampled in three sites across a proximal soil transect at seven dates over an 18 months period representing contrasted flooding periods in the wetland and we applied amplicon sequencing of the bacterial 16S rRNA gene and viral g23 gene (MiSeq Illumina).

## 2. Materials and methods

### 2.1. Study area, soil samplings and soil properties

Soil samples were collected from a coastal freshwater wetland classified as Natura 2000 and located near the city of Guidel (Brittany, Western France). It is part of the French network of hydrogeological observatories (H+ hplus.ore.fr/en), the OZCAR Research Infrastructure (Gaillardet *et al*., 2018) and the Socio-Ecological (eLTER) observatories network. Three closely spaced soil sampling sites (MP2, MP3 and MP4), located 3 to 10 meters apart, were selected based on the position of instrumented piezometers already installed on the wetlands. These sites exhibited distinct vegetation cover, MP2 and MP3 were situated beneath oaks trees, whereas MP4 was dominated by *Phragmites australis*. Soil samples were collected from each site at seven sampling dates from January 2022 to June 2023 (24 January, 18 May, 16 September, 12 November in 2022 and 29 January, 17 March, 12 June in 2023). Soil samplings were performed at 20 cm around each piezometer using a soil auger on the first 15 cm of depth. For each sampling point, three soil replicates were pooled together in sterile plastic bags and transported to the laboratory at 4°C. Soil samples were passed through a 2 mm mesh sieve and divided in subsamples for subsequent analyses. Five grams of soil (n=3) were used to determine the soil water content and 0.5g of soil (n = 4 to 6) were weighted in Lysing matrix E tubes (MP Biomedicals, USA) and stored at −20°C for DNA extraction. Soil properties (texture, water pH, cationic exchange capacity (CEC), contents in organic matter (OM), organic carbon (SOC) and total Nitrogen) from the three sampling sites (MP2, MP3 and MP4) were determined at the beginning of the sampling campaign on the soil sampled on January 2022 by Aurea Agrosciences laboratory (Ardon, France-http://www.aurea.eu).

### 2.2. Soil DNA extraction, PCR amplification and sequencing

DNA was extracted according to the protocol of Griffiths (2000), adapted by Nicolaisen et al.(2008) and Monard et al. (2016). The DNA quality was assessed on a 1% agarose gel and spectrophotometrically on a SpectroStarNano plate spectrophotometer (BMG Labtech, Germany).The V3-V4 regions of the 16S rRNA gene were amplified with the 341F-785R primers (Klindworth *et al*., 2013) and the major capsid gene, g23 of T4-type phages was amplified with the primers MZIA 1bis (5′-GAT ATT TGI GGI GTT CAG CCI ATG A-3′) and MZIA6 (5′-CGC GGT TGA TTT CCA GCA TGA TTT C-3′) (Dar *et al*., 2023). Each primer set contained the following overhang adapters: forward overhang R1 (5′-ACA CTC TTT CCC TAC ACG ACG CTC TTC CGA TCT-3′) and reverse overhang R2 (5′-GTG ACT GGA GTT CAG ACG TGT GCT CTT CCG ATC T-3′). PCRs were carried out in two replicates in a total volume of 25 μL containing each 0.4 µM primers and 0.8 mM dNTPs and 2 µL of 10-fold diluted DNA. For 16S rRNA gene amplification, the reaction was performed with 12.5 µL 2X Phanta Max Super Fidelity buffer (Vazyme, China), 1 µL Phanta Max Super-Fidelity DNA Polymerase (5U; Vazyme, China) and ultrapure water to reach the final volume. The amplification conditions were as follows: 3 min at 95°C, 30 cycles of 15 s at 95°C, 30 s at 65°C and 30 s at 72 °C and a final 5 min extension step at 72 °C. For g23 gene amplification, the PCR mixture contained 2.5 µL 10X TransTaq HiFi buffer (TransGen Biotech, China), 0.5 µL TransTaq-T DNA Polymerase (2.5 U; TransGen Biotech, China), 50 ng of T4 gene 32 protein (New England Biolabs, USA) and ultrapure water to reach the final volume of 25 µL. The amplification conditions were as follows: 3 min at 94°C, 35 cycles of 15 s at 94°C, 1 min at 54°C and 45 s at 72°C and a final 5 min extension step at 72 °C. The replicate amplicons were pooled and the PCR products targeted the g23 gene were separated using a 1.2% agarose gel electrophoresis. Bands corresponding to PCR products from 400 to 500 bp were cut and the sliced gel agarose was processed with the NucleoSpin Gel and PCR Clean-up kit (Macherey-Nager, The Netherlands) to purify PCR products from the expected size. 16S rRNA and g23 amplicons were sent to the Génome Québec Innovation Center for normalization, barcoding, multiplexing and Illumina paired-end (2 × 300 bp) MiSeq sequencing.

### 2.3. 16S rRNA gene quantification

Bacterial abundance was determined by quantitative PCR targeting the 16S rRNA gene on a LightCycler480 (Roche Diagnostics, Austria) and using the 341F-785R primers (Klindworth *et al*., 2013). Reaction mixtures (12.4 μl) were composed of 6 μl SensiFAST SYBR No-ROX mix (Bioline, England), 0.4 μl of each primer (10 μM), 4.6 μl ultrapure water and 1μl100-fold diluted DNA template. The qPCR program consisted in 5 min at 95°C, followed by 40 cycles of 15 s at 95°C, 30 s at 60°C, 30 s at 72°C and a final melting curves step consisting in 15 sec at 95°C, 1 min at 65°C with an increase of 0.5°C/5 s from 65°C to 95°C. Standard curves were generated by serial dilutions of genomic DNA of *Pseudomonas fluorescens* SBW25 to obtain 16S rRNA gene copy numbers ranging from 10^2^ to 10^6^. PCR efficiency was of 80 %. Potential inhibition was assessed by quantification of the 16S rRNA gene copy numbers in serial dilutions of a selection of soil DNA samples leading to the choice of 100-fold diluted DNA template for the qPCR. The quantities 16S rRNA gene fragments were expressed in gene copy numbers per g of dry soil.

### 2.4. Bacterial and viral community analyses

Bacterial and viral g23 de-multiplexed raw fastq files were processed with the DADA2 pipeline (Callahan *et al*., 2016) on Galaxy platform (https://usegalaxy.eu/). All primers were removed from each sequence using Cutadapt (Martin, 2011) and merged reads from 375 to 500 bp were selected for bacterial dataset whereas no size selection was applied for g23 dataset. Bacterial and viral amplicon sequence variants (bASVs and vASVs, respectievely) were inferred using the default pipeline in DADA2 and SILVA version 138 (Yilmaz *et al*., 2014) was used for bacterial taxonomic assignation. Sequences were deposited to the European Nucleotide Archive under project number PRJEB98874. Downstream analyses were conducted using RStudio (v2024.04.02) and R version 4.4.1. The Phyloseq package (McMurdie and Holmes, 2013) was used to remove ASVs not assigned to bacteria in the bacterial dataset, ASVs with less than 5 sequences and those detected in fewer than 3 independent samples. Metacoder (Foster *et al*., 2017) was used to visualize bacterial community compositions. Amino acid sequences of viral ASVs (vASVs) were obtained using getorf (Rice *et al*., 2000; Olson, 2002) with a minimum ORF size of 300 nucleotides and sequence matches were determined using BLASTp with the Diamond alignment tool (Buchfink *et al*., 2014) in Galaxy. vASVs with amino acid sequences coding for g23 capsid gene were selected and CD-HIT (Fu *et al*., 2012) was run with a threshold of one to remove redundant ASVs based on their amino acid sequences. A total of 165 vASVs were recovered.

Prior to alpha-diversity analyses, bacterial and viral datasets were rarefied to 10 035 and 799 sequences per sample, respectively, using the rarefy_even_depth function of the Phyloseq package. Alpha diversity for both bacterial and viral ASVs were assessed using the Shannon diversity index, as this diversity measure accounts for both richness and evenness within each sample.

Bacterial ASV (bASV) abundances were normalized based on the 16S rRNA gene copy numbers quantified by qPCR. Beta diversity of both bacterial and g23 ASV communities was assessed via Bray-curtis dissimilarity metrics, calculated with the ordinate and distance functions from the phyloseq package. Results were plotted by performing (PCoA) with the function plot_ordination and ellipses (95% confidence) highlighting clustering of sample- and date-specific communities were drawn using the function stat_ellipse of the ggplot2 package (Wickham, 2016). Differences in the Bray−Curtis distance metrics over time and between sites (MP2, MP3 and MP4) were analyzed using the Vegan package (Oksanen *et al*., 2015)with PERMANOVA (adonis2 function) and community dispersion was estimated with the function betadisper. Venn diagrams were constructed using the trans_venn function of the microeco package (Liu *et al*., 2021). A heatmap was generated (pheatmap package) on the 40 most abundant viral ASVs after normalization by log-transformation and ASVs were hierarchically clustered using Euclidean distance and complete linkage.

### 2.5. T4-type phage phylogenetic analysis

Two clustering steps were performed. First, amino acid sequences of g23 were clustered using CD-HIT (Fu *et al*., 2012), representative sequences were established at 70% and their closest relative were identified using BLASTp on Diamond alignment tool (Buchfink *et al*., 2014) in Galaxy. These close relative sequences, sequences from two *Escherichia coli* T4 phage isolates (Subedi and Barr, 2021) and 113 amino acid sequences of g23 sequenced from different environments (Antarctic glacier (Bellas and Anesio, 2013), freshwater (López-Bueno *et al*., 2009), pond freshwater in USA (Dar *et al*., 2023), diverse marine open waters (Filée *et al*., 2005), rice paddy soils in Japan (Fujii *et al*., 2008; Wang *et al*., 2009a; Cahyani *et al*., 2009) and China (Wang *et al*., 2009a, 2009b; Liu *et al*., 2012; Li *et al*., 2019a), rice paddy water in China (Liu *et al*., 2016), wetlands soil and water (Zheng *et al*., 2013), upland soil in China (Wang *et al*., 2011; Liu *et al*., 2011), subtropical estuary sediments (He *et al*., 2017), dairy wastewater (WW) waters in United States (Jamindar *et al*., 2012) and China (Liu *et al*., 2017)) were downloaded from GenBank (NCBI) and merged with the sequences generated in this study. Then, the entire g23 dataset was clustered with CD-HIT (Fu *et al*., 2012) with a threshold of 0.7. Clustered sequences were aligned (complete alignment) with ClustalW on Galaxy and maximum likelyhood tree was built using PHYLIP and PhyML (Anisimova and Gascuel, 2006; Dereeper *et al*., 2008; Guindon *et al*., 2010) with default settings, respectively. The unrooted tree was visualized using the Interactive Tree of life online software (Letunic and Bork, 2024).

### 2.6. Network analysis

To construct co-occurrence networks, bacterial and g23 ASV tables were merged and co-occurrence analysis was performed on the entire dataset using the sparCC method (Friedman and Alm, 2012) implemented in the microeco package. ASVs with a relative abundance greater than 0.1% (filter_thres) were retained for network construction (Liu *et al*., 2021). Eigen gene values of each module from the co-occurrence network of the entire dataset and involving at least three nodes were used for correlation analysis with soil moisture using the functions cal_eigen and cal_cor. The network was built with the cal_network function and visualized using Gephi (Bastian *et al*., 2009). Each node represented one bacterial or viral ASV, and edges connecting the nodes represented significant positive or negative Pearson correlations (*p*< 0.01) between two ASVs.

### 2.7. Statistical analyses

Significant differences in the initial soil properties, in soil moisture, in viral Shannon diversity index and in 16S gene copy numbers between sampling sites were tested using Kruskal-Wallis rank test (kruskal function from the agricolae package). Significant differences in the bacterial Shannon diversity index between sampling sites was tested using one-way ANOVA (aov function) followed by Tukey’s HSD test. Correlations between soil moisture contents, 16S gene copy numbers and diversity indexes were calculated using Kendall Tau coefficient (cor.test function from stats package). Repeated-measures ANOVAs were performed to assess the effects of sampling date, site, and their interaction on diversity indexes and relative abundance of bacterial classes using the aov function in the stats package in R.

## 3. Results

### 3.1. Bacterial and viral diversity across sites

Soil properties of the three sites are presented in Table 1. A gradient in texture was observed with sand content decreasing and clay content increasing from MP2 through MP3 to MP4 (Kruskal–Wallis test, *p* = 0.03 for both). Total nitrogen content also increased significantly along this gradient (Kruskal–Wallis test, *p* = 0.02) while no significant differences were observed for both OM and SOC contents (Kruskal–Wallis test, *p* = 0.08 for both) leading to a decreasing C/N ratio from MP2 to MP4.

**Table 1.** Soil properties of the three sampled sites.

|  | MP2 | MP3 | MP4 |
| --- | --- | --- | --- |
| Sand (%) | 19.98 ± 0.19a | 13.57 ± 0.11b | 6.73 ± 0.40c |
| Loam (%) | 49.26 ± 0.97a | 51.52 ± 1.29a | 51.47 ± 3.58a |
| Clay (%) | 30.76 ± 1.16a | 34.90 ± 1.18b | 41.80 ± 3.18c |
| OM (%) | 8.77 ± 0.40a | 8.90 ± 1.31a | 9.90 ± 0.10a |
| SOC (%) | 5.10 ± 0.25a | 5.18 ± 0.77a | 5.77 ± 0.05a |
| Total N (%) | 0.51 ± 0.01a | 0.58 ± 0.01b | 0.72 ± 0.01c |
| pH<sub>w</sub> | 5.93 ± 0.06a | 6.10 ± 0.10a | 5.33 ± 0.06b |
| CEC (meq/100 g) | 29.20 ± 1.06a | 30.37 ± 1.81ab | 34.40 ± 1.18b |

At each sampling date, soil moisture content was also measured, revealing a significant gradient across the three sites (Fig. 1D) on top of the seasonal fluctuations (Fig. S1). MP4 was the wettest site with a mean moisture content of 179.54% ± 57.08 while MP2 was the driest with 80.89% ± 27.26 (Fig. 1C). Bacterial abundance was significantly lower in MP4 compared to the two other sites (Fig. 1D, Kruskal-Wallis test, *p* = 0.002) and the bacterial 16S rRNA gene copy number was significantly inversely correlated to the soil moisture (Kendall’s tau rank correlation, R^2^ = −0.22, *p* = 0.003).

**Fig. 1.**
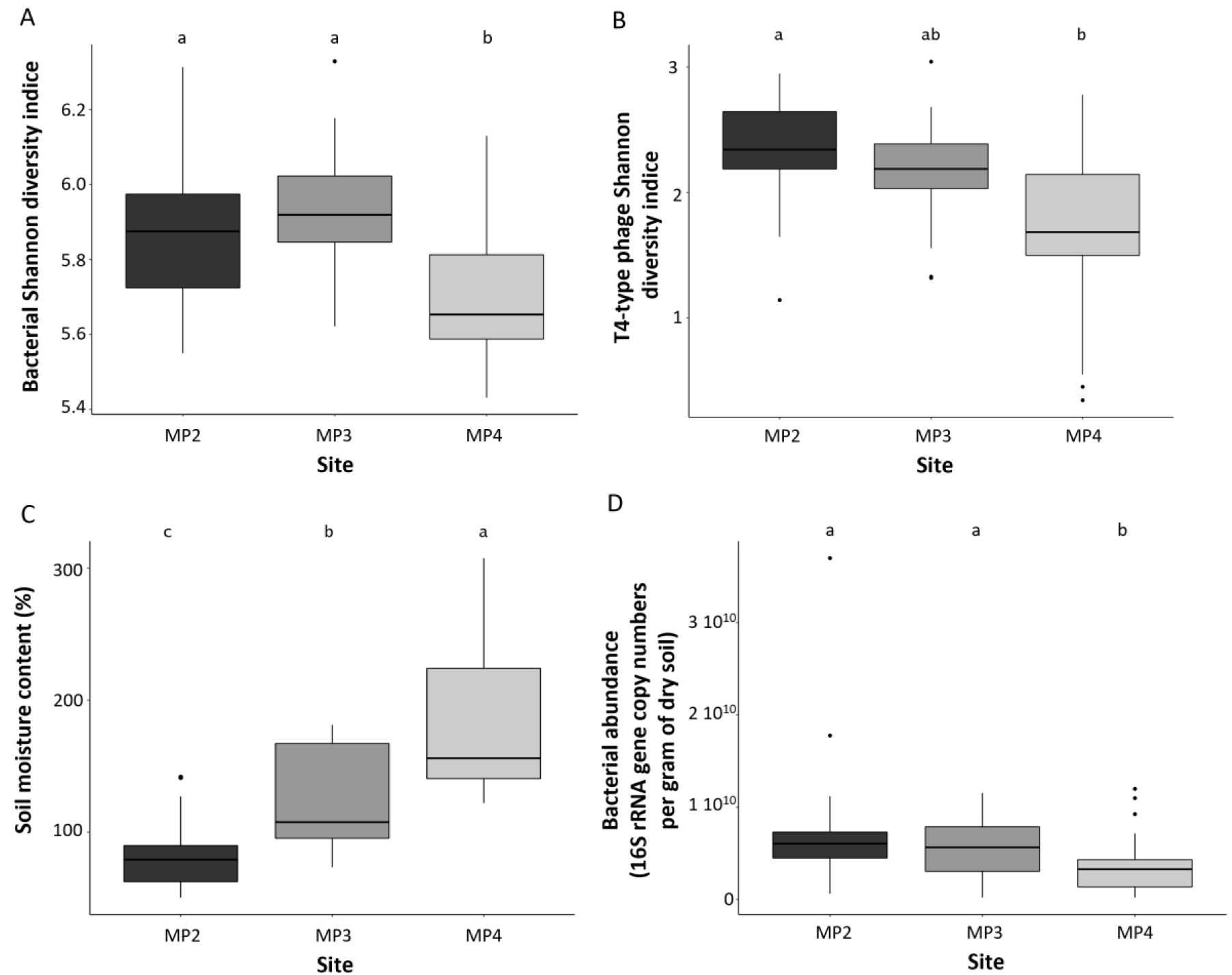
Bacterial (A) and T4-type phages (B) Shannon diversity indexes, soil moisture content (C) and bacterial abundance (D) in the soils from three sampling sites (MP2, MP3 and MP4). In each box plot, the top and bottom of each box represent the 25th and 75th percentiles, and the center line indicates the median. Different letters indicate statistically significant differences at *P* <0.05 between the sites.

The dynamics of bacterial and viral Shannon diversity indexes were investigated with time in the three sampling sites. A site (2-ways ANOVA, F = 5.656, *p* = 0.034; F = 50.150, *p* = 0.019, respectively) and date effects (2-ways ANOVA, F = 4.157, *p* = 0.002; F = 7.567, *p* = 4.1710^-5^, respectively) were identified for both, as well as an interaction effect on viral alpha diversity (F = 2.968, *p* = 0.007; Fig. S2). Independently of the sampling date, changes in the diversities of bacteria and T4-type phages were also observed between the three sites, the Shannon diversity indices being significantly lower in MP4 than in MP2 and MP3 for bacteria (One-way ANOVA, F = 11.16, *p* = 5.25 10^-5^) and in MP2 for T4-type phages (Kruskal-Wallis, test *p* = 0.001), respectively (Fig.1A and B).

### 3.2. Bacterial communities are primarily structured across sites

Whatever the sampling site, the bacterial communities were dominated by the following classes: *Planctomycetes, Vicinamibacteria* (mainly belonging to the *Vicinamibacterales* order), *Methylomirabilia* (mainly belonging to the *Rokubacteriales* order), *Verrucomicrobiae,* KD4-96, *Alphaproteobacteria*, *Phycisphaerae (*mainly belonging to the *Tepidisphaerales* order), *Thermoleophilia* and *Gammaproteobacteria* (mainly belonging to the *Burkholderiales* order) (Fig.2). A core microbiota was detected in the three sites representing 77.9% of the bacterial sequences (Fig. S3A). Besides these common taxa, bacterial communities were significantly different between sites (Adonis test R^2^ = 0.32, *p* = 0.001) and depending on the sampling dates (Adonis test R^2^ = 0.17, *p* = 0.001) (Fig. 3A). Among the most abundant bacterial classes, site-related differences were primarily driven by significantly lower relative abundances of *Phycisphaerae*, *Planctomycetes* and *Vicinamibacteria* in MP4 compared to the other sites (2-ways ANOVA; F = 15.162, *p* = 2.86 10^-3^; F = 10.227, *p* = 8.37 10^-3^; F = 41.385, *p* = 1.32 10^-4^, respectively), and by significantly higher relative abundances of *Gammaproteobacteria* and *Methylomirabilia* in MP4 (2-ways ANOVA, F = 40.112, *p* = 1.46 10^-4^; F = 171.891, *p* = 1.12 10^-6^, respectively) (Fig. 4). Linked to the moisture gradient observed across the three sampling sites (Fig. 1C), the relative abundances of these bacterial classes were significantly positively (*Gammaproteobacteria* R^2^ = 0.24, *p* = 0.001 and *Methylomirabilia* R^2^ = 0.15, *p* = 0.043) or negatively (*Phicisphaerae* R^2^ = −0.21, *p* = 0.006*, Planctomycetes* R^2^ = −0.18, *p* = 0.015 and *Vicinamibacteria* R^2^ = −0.28, *p* = 2.53 10^-4^) correlated to soil water content, based on Kendall’s tau rank correlation.

**Fig. 2.**
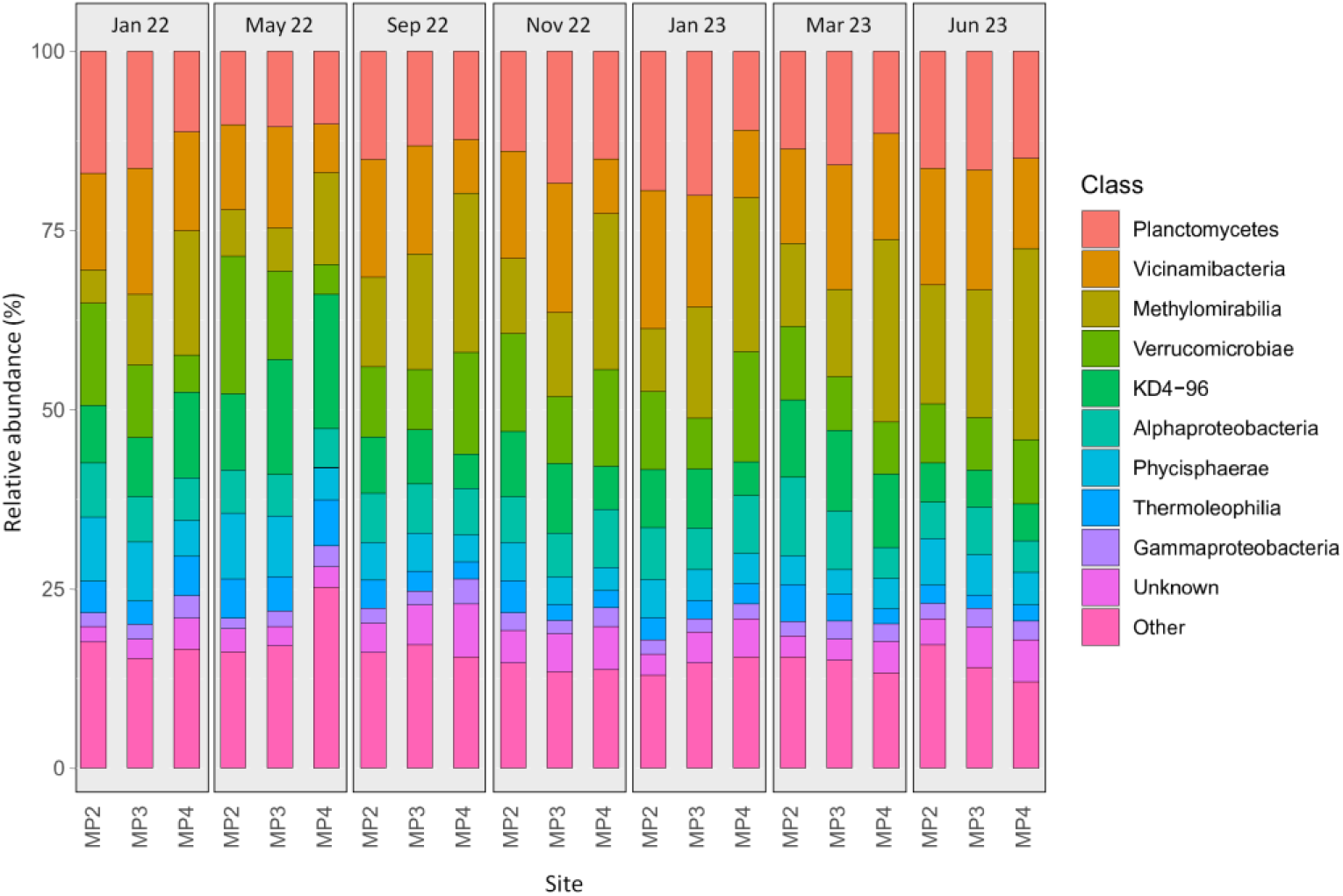
Relative abundance of the main bacterial classes in the three sites (MP2, MP3, MP4) at the seven sampling dates (January 2022, May 2022, September 2022, November 2022, January 2023, March 2023 and June 2023).

**Fig. 3.**
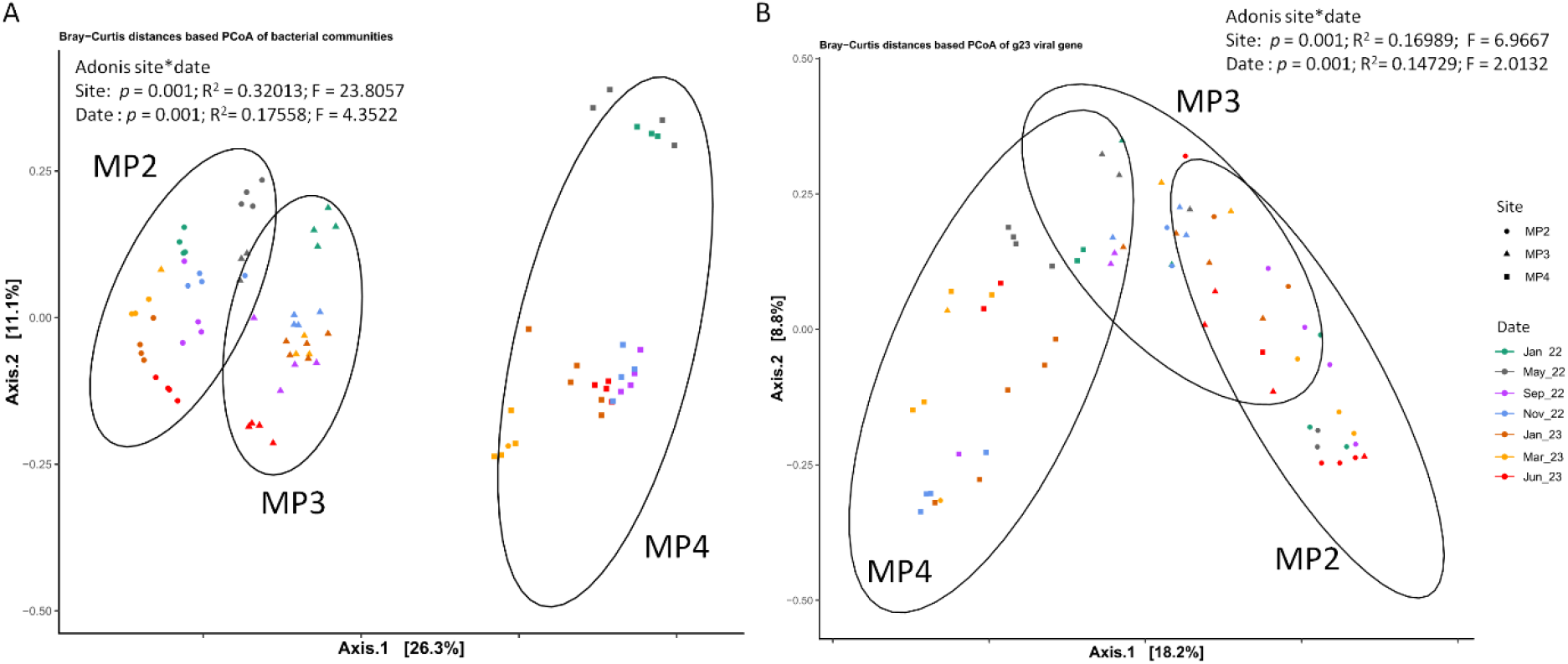
Community structure for bacteria (A) and T4-type phages (B) - Principal coordinates analysis (PCoA) based on Bray-Curtis distance of bacterial and viral communities in the three sampling sites over time. Different colors represent different sampling times and different shapes represent different sites. *F, R^2^*and *P* values are given (based on PerMANOVA). Ellipses represent 95% confidence intervals.

**Fig. 4.**
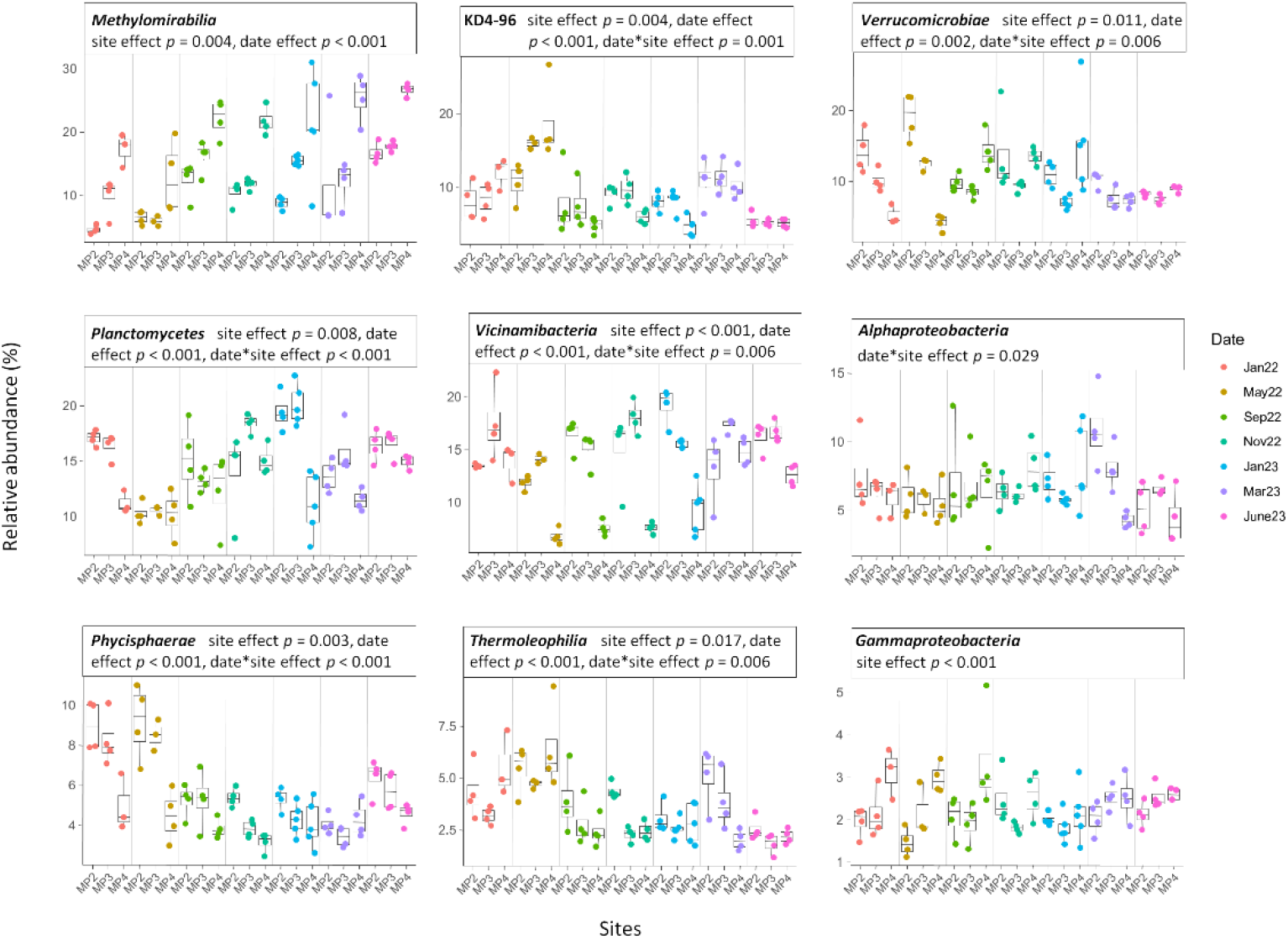
Changes in the relative abundance of the main bacterial classes in the three sites with time. In each box plot, the top and bottom of each box represent the 25th and 75th percentiles, and the center line indicates the median. Significant effect of date, site and their interactions are presented (Repeated-measures ANOVAs).

A temporal dynamic of the relative abundance of some bacterial classes was observed with significant changes in the proportions of KD4-96 (2-ways ANOVA, F= 19.448, *p* = 9.63 10^-12^), *Methylomirabilia* (2-ways ANOVA, F = 10.74, *p* = 9.75 10^-8^), *Phycisphaerae* (2-ways ANOVA, F = 28.434, *p* = 8.64 10^-15^), *Planctomycetes* (2-ways ANOVA, F=15.72, *p* = 3.4210^-10^), *Thermoleophilia* (2-ways ANOVA, F=13.628, *p* = 3.17 10^-9^), *Verrucomicrobiae* (2-ways ANOVA, F = 3.991, *p* = 2.33 10^-3^) and *Vicinamibacteria* (2-ways ANOVA, F = 10.110, *p* = 2.18 10^-7^) depending on sampling dates (Fig. 4). At the ASV level, the MP4 site also stood out from the others, hosting the highest proportion of site specific bASVs with 457 bASVs corresponding to 2.3 % of the total number of bacterial sequences, compared to 217 and 174 bASVs specific to MP2 and MP3, respectively (Fig. S3A). Additionally, while 782 ASVs were shared between MP2 and MP3, only 338 and 517 were common between MP4 and MP2, and between MP4 and MP3, respectively (Fig. S3A).

### 3.3. Viral communities

As observed for bacteria, viral communities were significantly different between sites (Adonis test R^2^ = 0.17, *p* = 0.001) and depending on the sampling dates (Adonis test R^2^ = 0.15, *p* = 0.001) (Fig. 3B). Sixty-nine out of the 165 viral ASVs, accounting for 74.4% of the total viral sequences, were detected across all three sites (Fig. S3B). Among these, vASV_1 was particularly abundant in MP2 and MP3, contributing for 10 and 5% of the total viral sequences at each site, respectively. When comparing the sites, MP2 and MP3 shared the highest number of viral ASVs (36), but they accounted for only 6.8% of the total viral sequences. In contrast, the 32 vASVs shared between MP3 and MP4 represented 16.3% of the total viral sequences (Fig. S3B). This greater sequence overlaps between MP3 and MP4 was largely driven by the high relative abundance of two viral ASVs in MP4, vASV_3 and vASV_5, which accounted for 6% and 4% of the total viral sequences at that site, respectively. The proportions of shared viral ASVs between sites were dynamic with time with the highest amount of common ASVs in the three sites observed in June 2023 (23 vASVs - 63.6 % of the total amount of sequences) and the lowest in November 2022 (2 vASVs - 2.5 % of the total amount of sequences; Table S1). Within the 40 most abundant g23 ASVs, spatial patterns were clearly observed (Fig. S4) with clusters of viral ASVs dominating in MP2 and in MP4.

Phylogenetic analysis of the 165 amino acid g23 sequences obtained in this study incorporated 115 reference sequences from a range of diverse environments (Fig. 5) as well as two sequences from isolates of *E. coli* T4 phages. These reference sequences did not cluster neither according to their localization (China, Japan, Antarctic, United States) nor to their environment (soil, fresh and marine waters, sediments, ice). Most of the g23 ASVs from the Guidel wetland soils clustered with known sequences previously obtained from diverse environments. However, some clades corresponding to 26 vASVs did not cluster with any of the reference sequences included in the phylogenetic analysis (branches labeled in red in Fig.5). Among them, the clade comprising vASV_35 and vASV_135 and the one with vASV_5, vASV_13 and vASV_42 were predominantly represented in MP3 and MP4, respectively.

**Fig. 5.**
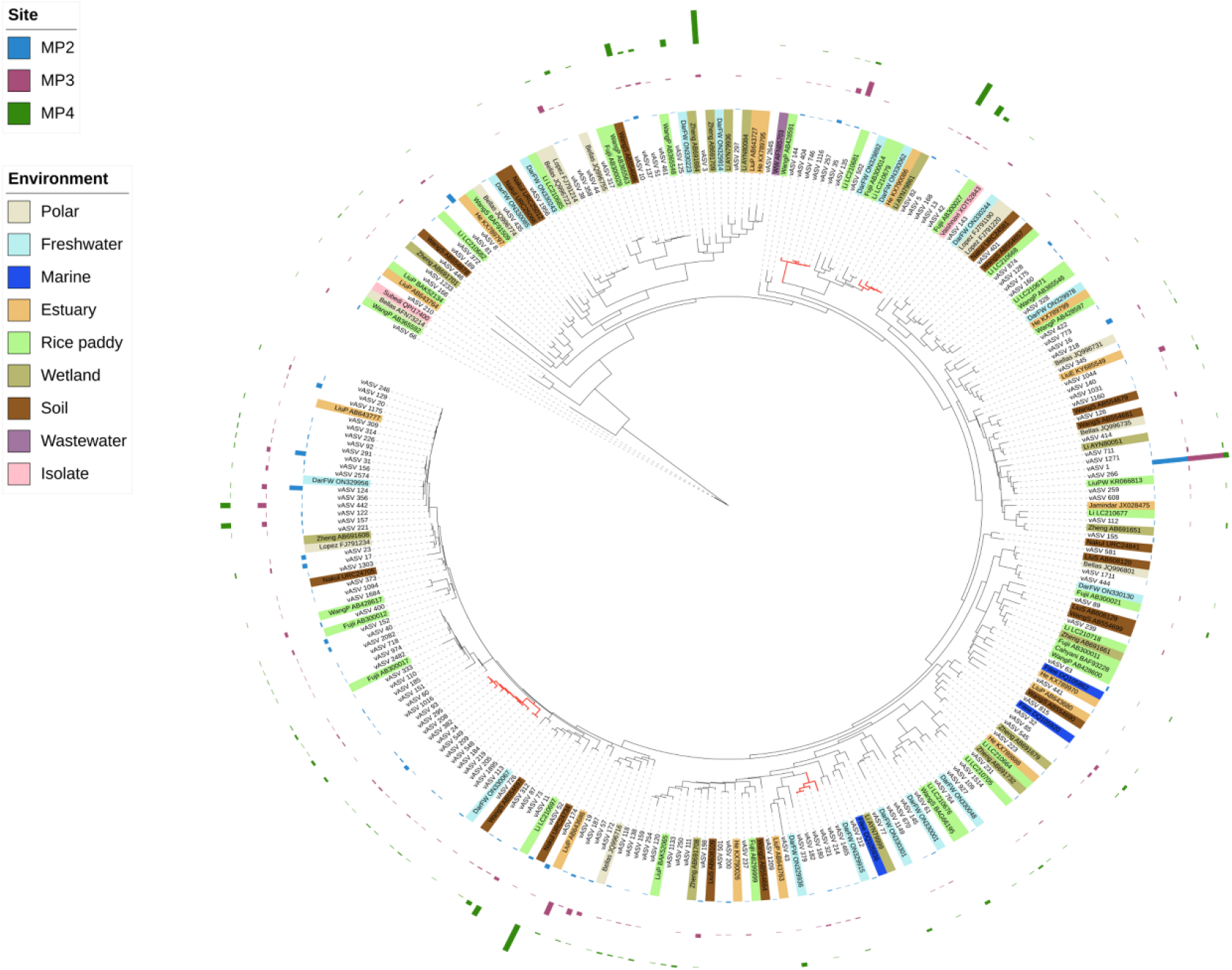
Unrooted maximum likelyhood phylogenetic tree based on 281 g23 amino acid sequences. Different colored ranges indicate sequences originating from diverse environments. vASV obtained in the present study are not colored. Branches of clades that did not cluster with any of the reference sequences are colored in red. The outer colored rings indicate the relative abundance of sequences of each vASV in each site.

The vASV_66 branched apart all the other sequences at the root of the tree (bootstrap values = 0.999) as two references sequences obtained from Antarctic glacier and Chinese rice paddy soil (Fig. 5; Bellas_AFN73214 and WangP_AB365592, respectively). While one of the two T4 phage isolate (Subedi_QPI17400) also branched apart at the root of the tree, the other sequence (Vaishnavi_XOT52843) clustered with vASV_143 (Fig. 5).

### 3.4. Co-occurrence network analysis

Co-occurrence network was constructed to explore the interaction patterns between bacteria and T4-type phages across the entire dataset (Fig. 6). It consisted in 83 nodes (50 bacterial and 33 viral) and 77 edges, among which 59 were positive. Correlations between bacterial ASVs accounted for 46.8 %, between viral ASVs for 5.2 % and between bacterial and viral ASVs for 48.0 %. While no negative association was observed between viral ASVs, 10 were detected between a bacterial and a viral ASV (Fig. 6A). These negative co-occurrence links involved bacterial ASVs belonging to various classes: KD4-96, *Vicinamibacteria*, *Methylomirabilia*, *Planctomycetes* and *Alphaproteobacteria*.

**Fig. 6.**
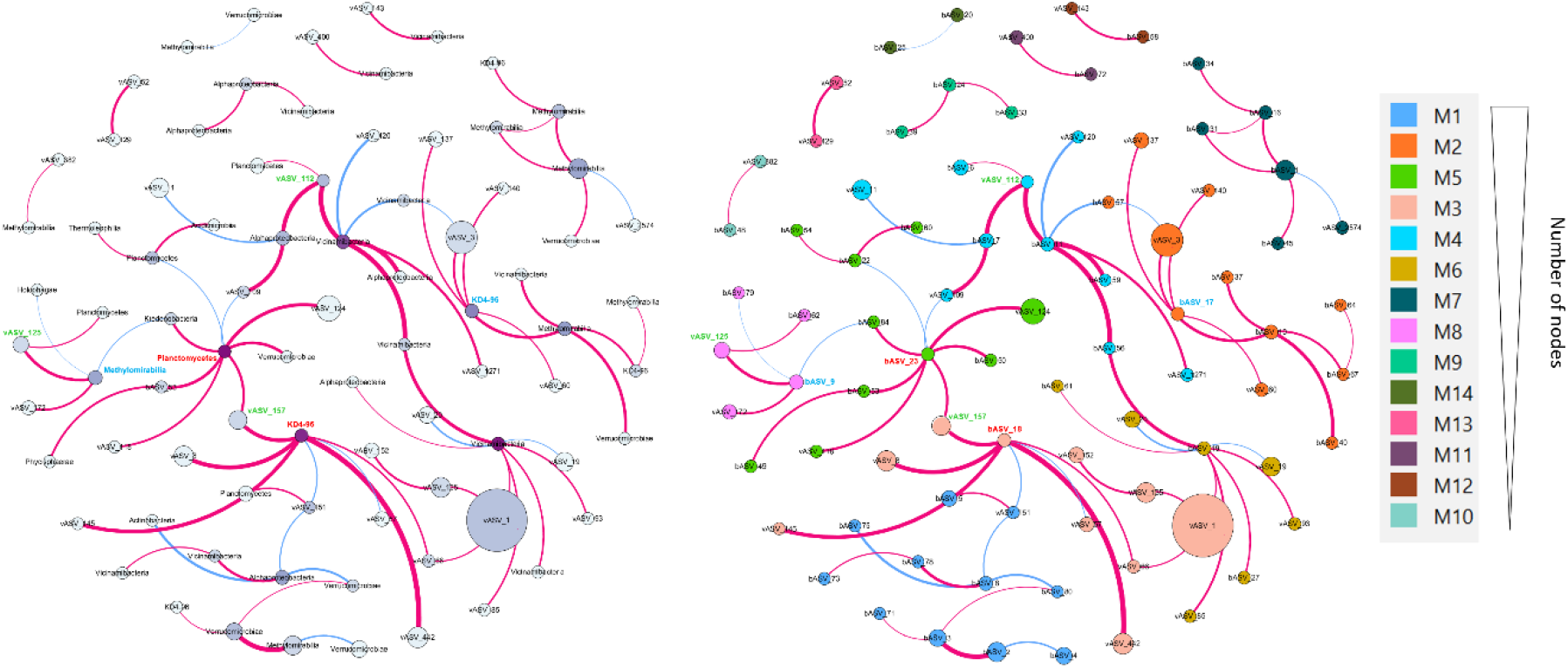
Co-occurrence network of bacterial and viral ASVs with either ASV and bacterial class (A) or module (B) labels. Blue and red lines represent negative and positive interactions with strong (r > 0.7 or r < −0.7) and significant Pearson correlations(*P*<0.01). Line width indicates the strength of the association (r) and the size of each node is proportional to its link numbers.

We identified 14 distinct modules, each representing areas of dense connections with sparser links with other areas (Fig. 6B). Within these modules, two bacterial nodes (bASV_18 - KD4-96 and bASV_23 - *Aquisphaera* sp.; labeled in red in Fig. 6) were identified as hubs, exhibiting high within-module connectivity (Z > 2.5). The modules M2 and M3 were significantly negatively correlated to soil moisture content (R^2^ = −0.40, *p* = 0.009; R^2^ = −0.35, *p* = 0.023, respectively) while the module M5 was significantly positively correlated to soil moisture content (R^2^ = 0.33, *p* = 0.030).

Seven bacterial ASVs were correlated to several viral ASVs and only bASV_9 (*Rokubacteriales* order; labeled in blue in Fig. 6) and bASV_17 (KD4-96; labeled in blue in Fig. 6) were always positively associated to viral ASVs. The other five bASVs were both positively or negatively correlated with multiple vASVs. Reciprocally, six viral ASVs were connected to two or three different bASVs and these bASVs belonged to several bacterial phyla. Among these six viral ASVs, three (vASVs_112, vASV_125 and vASV_157; labeled in green in Fig. 6)) were always positively associated to their connected bASVs.

## 4. Discussion

Wetlands are particular habitats for soil microorganisms, characterized by fluctuations in water table levels inducing successions of anoxic and aerobic conditions in presence or absence of water saturation, respectively. These changes in the hydrological regime affect the geochemistry of wetland soils through induced shifts in microbial community structures (Unger *et al*., 2009; Nunes *et al*., 2015), leading to slower SOM decomposition and specific microbial assemblages in these habitats. In the present study, we investigated both bacterial and T4-type phage communities in a wetland soil across three sites gradually influenced by the water table level, MP4 being the site presenting the highest moisture content.

### 4.1. Site-specific shifts in bacterial community composition and potential functions

The dominant bacterial communities were similar across the three sites, with a significant proportion of bASVs belonging to a shared core microbiota. However, we identified two bacterial classes, *Methylomirabilia* and *Gammaproteobacteria*, which were significantly enriched in MP4 compared to the two other sites. Within *Methylomirabilia, Rokubacteriales* was the most representative order and these bacteria have previously been detected in wetland soils (e.g. Ivanova *et al*., 2022). While the physiology of these bacteria is poorly understood, they have been identified as neutrophilic, non-methanotrophic bacteria (Ivanova *et al*., 2022; Lan *et al*., 2022; Kang *et al*., 2024). As previously observed (Ivanova *et al*., 2022), their relative abundance increased with the N content which was the highest in MP4. 13.5% of the *Methylomirabilia* bASVs were affiliated to the *Methylomirabilales* order that are methanotrophic bacteria (Peres *et al*., 2023). The *Gammaproteobacteria* were dominated by bASVs from the *Burkholderiales* order, some members of which play important role in nitrogen cycle through denitrification and N_2_ fixation (Tang *et al*., 2019; Li *et al*., 2019b). Ten percent of the *Burkholderiales* bASVs belonged to the *Gallionellaceae* family which are microaerobic iron-oxidizing bacteria (FeOB) playing major role in the Fe biogeochemical cycle in freshwater (Hedrich *et al*., 2011). These FeOB have been previously detected in abundance in the anoxic, Fe-rich groundwater which feeds the Guidel wetland (Bethencourt *et al*., 2020).

The relative abundances of some of the dominant bacterial classes (*Phycisphaerae*, *Planctomycetes* and *Vicinamibacteria*) were significantly reduced in MP4 compared to MP2 and MP3 and these bacteria are known to contribute to overall carbon cycling. Members of the *Tepidisphaerales* order that dominated the *Phycisphaerae* class have been found to possess a high number of carbohydrate-active enzyme (CAZyme) gene clusters highlighting their potential for complex carbon degradation (Lenferink *et al*., 2024). *Planctomycetes* genomes encode wide repertoires of CAZymes conferring them the ability to grow on a wide diversity of substrates such as xylan, pectin, starch, lichenan, cellulose, chitin and polysaccharides of microbial origin (Dedysh and Ivanova, 2019). Finally, members of *Vicinamibacterales* order that dominated the *Vicinamibacteria* class are recognized to be plant growth-promoting rhizobacteria (PGPR) (Wu *et al*., 2021) and they are involved in heterotrophic respiration in wetlands following low molecular weight compounds inputs (Zhang *et al*., 2021; Wang *et al*., 2023).

The significant differences in the relative abundance of bacterial classes in MP4 compared to the other two sites support our first hypothesis that bacterial community composition varies across a proximal soil transect, even within close habitats. Furthermore, given the central role of these dominant bacterial taxa in both nitrogen and carbon cyclings, their interactions with phages could influence soil biogeochemical processes.

### 4.2. A large diversity of T4-type phages was recovered in wetland soils

T4-type phages are widely distributed in diverse natural environments (Desplats and Krisch, 2003; Filée *et al*., 2005; López-Bueno *et al*., 2009; Fujihara *et al*., 2010; Li *et al*., 2019a) and present a large genetic diversity (Dar *et al*., 2023). This high genetic diversity was also observed at our study scale with vASVs clustering with reference sequences obtained from various localizations (China, Japan, Antarctic, United States) and environments (soil, fresh and marine waters, sediments, ice), consistent with the findings reported by Dar et al. (2023). The high similarity between g23 sequences obtained from different biomes could be explained by either the presence of identical bacterial hosts in the different environments or more likely by a broad host range phages that are widely distributed in natural environments (Breitbart and Rohwer, 2005; de Jonge *et al*., 2019). However, four novel T4-type phage clades were also detected, with g23 sequences that did not cluster with any previously described sequences.

While a large diversity of T4-type phages was observed across our three sampling sites, the viral community was dominated by few vASVs, which were highly represented in one or two sites. This observation supports the Bank model proposed by Breitbart and Rohwer (2005), which states that only the most abundant viruses are active, whereas the majority are inactive, rare and constitute a ‘bank’ of viruses able to infect new hosts that would grow with changing environment. This is in line with the observations of Chow et al. (2014) and Casteldine and Buckling (2024), who reported that, at seasonal timescales, viruses may more often follow their hosts rather than control host abundance. In the present study, PCoA analysis suggests that both bacterial and viral communities exhibit niche partitioning. Given that both communities are similarly structured across the three sites, T4-type phages may be following bacterial population dynamics. Consequently, changes in the dominant host population are predicted to result in changes in the active viral fraction (Breitbart and Rohwer, 2005).

### 4.3. Investigation of host-phage preferential associations

T4-type phages are known to infect bacterial host from various genera of *Pseudomonadota* and *Cyanobacteria* (Nakayama *et al*., 2009; Petrov *et al*., 2010; Sullivan *et al*., 2010; Roux *et al*., 2015). In the present study, although one vASV clustered with the g23 sequence of the *E. coli* phage KJC1000, which was isolated from sewage water in India, no members of the order *Enterobacterales* was detected in our bacterial dataset. This suggests that closely related T4-type phages may have a broad host range. Moreover, despite the high diversity of vASVs detected, only the bacterial orders *Sphingomonadales* and *Aeromonadales*, both known to include hosts for T4-type phages, were detected in our dataset. These results imply that the majority of T4-type phage hosts is still unknown. Since many host bacteria remain uncultivable, interactions between environmental phages and their hosts remain poorly studied. As an alternative to laboratory techniques, co-occurrence networks relying on cultivation independent methods have been proposed to predict phage-host associations, with positive correlations suggesting similar preferred conditions and negative correlations indicating viral infection and lysis (Chow *et al*., 2014; Li *et al*., 2019a, 2021). Interpretation of such networks should be undertaken with caution given the several limitations identified when applied to soil microbial communities. First, co-occurrence networks do not reflect the effective interactions between soil microorganisms since these interactions rely mainly on microbial cells that individually respond to local conditions, independently of their taxonomy. Then, these networks are generally based on data obtained from soil samples that are not representative of the microscale at which microorganisms interact. Finally, ecological interpretation of co-occurrence network is limited by the fact they do not distinguish direct interactions between two ASVs from indirect ones mediated by a third ASV, or through bottom-up control when considering viruses and their hosts (Goberna and Verdú, 2022; Guseva *et al*., 2022). Considering these concerns, in the present study we applied co-occurrence networks to identify viral ASVs which tended to co-occur more frequently with bacterial ASVs, suggesting potential host-phage relationships. Such preferential associations were detected either between a single bacterial ASVs and several viral ASV or inversely, between a single viral ASV and several bacterial ASVs, suggesting either a susceptibility of host to multiple phages or a broad-host range for the T4-type phage considered. Interestingly, the co-occurrence network was dominated by positive correlations, which should reflect that viral ASVs are primarily controlled by host availability (Chow *et al*., 2014), confirming our previous observations based on the ‘Bank’ model with T4-type phages following bacterial population dynamics.

### 4.4. Soil moisture as a potential regulator of bacterial infection rates

The effects of changes in soil moisture contents were investigated in the present study through both temporal and spatial dynamics. Indeed, seasonal fluctuations in the water table levels induced significant changes in soil moisture with sampling dates (data not shown) and a significant gradient of soil moisture was observed across the three sampling sites. Among soil properties, moisture has previously been identified as a regulator of both microbial and viral life in soil (Jansson and Wu, 2022; Ma *et al*., 2024). Increased soil water content enhances connectivity between distant pores, potentially linking previously isolated microbial microniches which should affect the diversity of both soil bacteria and phages, as observed in MP4, the wettest soil in our study. Moreover, increases in soil moisture might improve encounter rates between phages and their hosts (Nicolas *et al*., 2022) and promote infection. Soil moisture also influences environmental ion concentration and ionic strength is critical for infection of bacteria by many phages (Carlson *et al*., 2023). Considering these key features of soil moisture, it is essential to broaden studies on wetland soil systems which by definition rely on this property. These soils contain more than 30% of the global total soil organic carbon (SOC) stock within 6% of the land surface (Poulter *et al*., 2021), and their status is currently threatened by climate change.

## 5. Conclusion

Soil phages are increasingly recognized for their important ecological roles, particularly through their influence on soil microbial dynamics and processes. This is expected to be especially significant in wetlands, which are widely acknowledged as some of the Earth’s most efficient ecosystems for carbon sequestration and storage. In the present study, we identified a large diversity of T4-type phages, among which a fraction was novel, while others were similar to phages previously sequenced from various biomes. These findings suggest that T4-type phages are capable of successfully colonizing diverse niches in the biosphere, contributing to their ubiquity and diversity. This ability may rely on their broad host range and their capacity to follow bacterial population dynamics. Our results provide new insights into the role of phages in soil, highlighting their interactions with bacterial hosts involved in carbon and nitrogen cycles, interactions that are likely regulated by fluctuations in soil moisture, as observed in wetlands.

## Acknowledgements

We would like to thanks Elisa Isola and Emmanuel Moutou for their help in the initial methodological developments and Dr Simon Chollet for his support in statistics. Genetic analyses were performed in the Molecular Ecology Platform PEM (UMR 6553 Ecobio, Rennes, CNRS/UR). This work was supported by the French National program EC2CO (Ecosphère Continentale et Côtière) from the CNRS-INSU and the Chaire ‘Biodiversity and climate change’ from the Fondation Université de Rennes.

## CRediT authorship contribution statement

**Rémi Trémouille**: Writing-review & editing, Acquisition of data, Analysis and interpretation of data.**Virginie Daburon**: Writing-review & editing, Methodology, Supervision, Investigation. **Achim Quaiser**: Writing-review & editing, Methodology, Conceptualization. **Alexis Dufresne**: Writing-review & editing, Methodology, Conceptualization. **Cécile Monard**: Writing-review & editing, Writing - original draft, Supervision, Resources, Project administration, Methodology, Funding acquisition, Conceptualization, Analysis and interpretation of data.

## Data availability

Amplicon sequencing data have been deposited to the European Nucleotide Archive under project number PRJEB98874.

## Supplementary data

**Supplementary Fig. 1.**
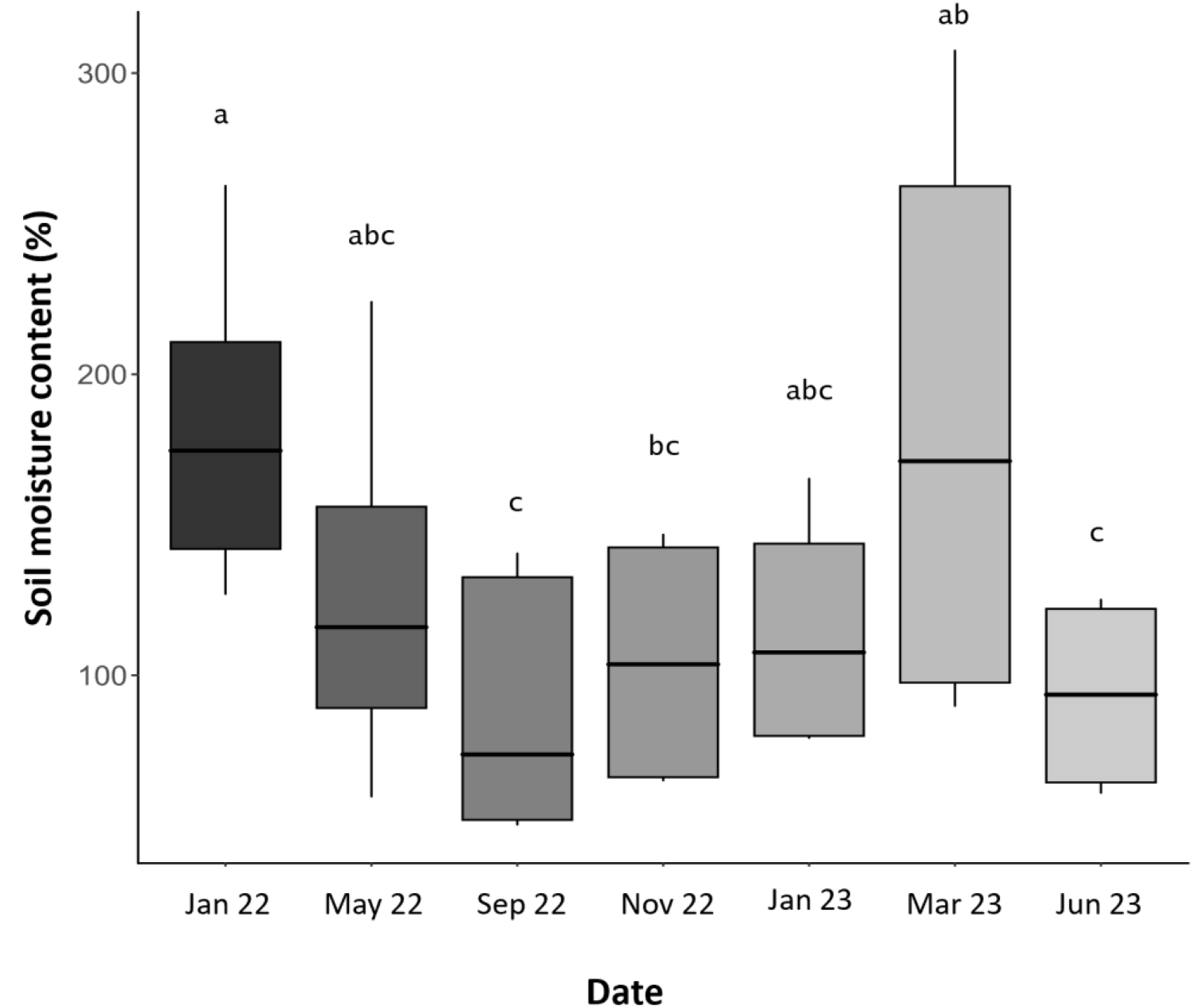
Mean soil moisture content across the three sampling sites at the different sampling dates.

**Supplementary Fig. 2.**
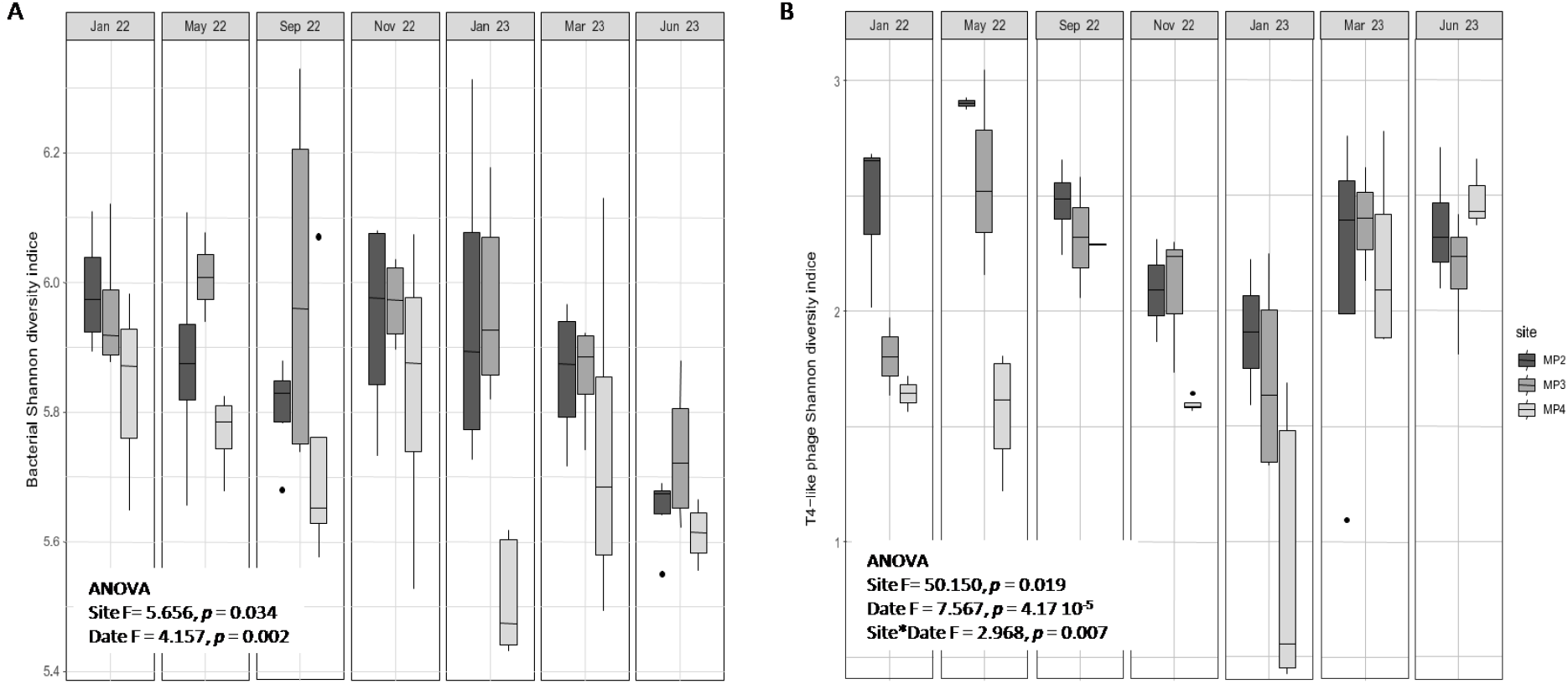
Bacterial (A) and T4-type phage (B) Shannon diversity indexes in the three sites at the seven sampling dates. In each box plot, the top and bottom of each box represent the 25th and 75th percentiles, and the center line indicates the median. Results of repeated-measures ANOVAs are presented.

**Supplementary Fig. 3.**
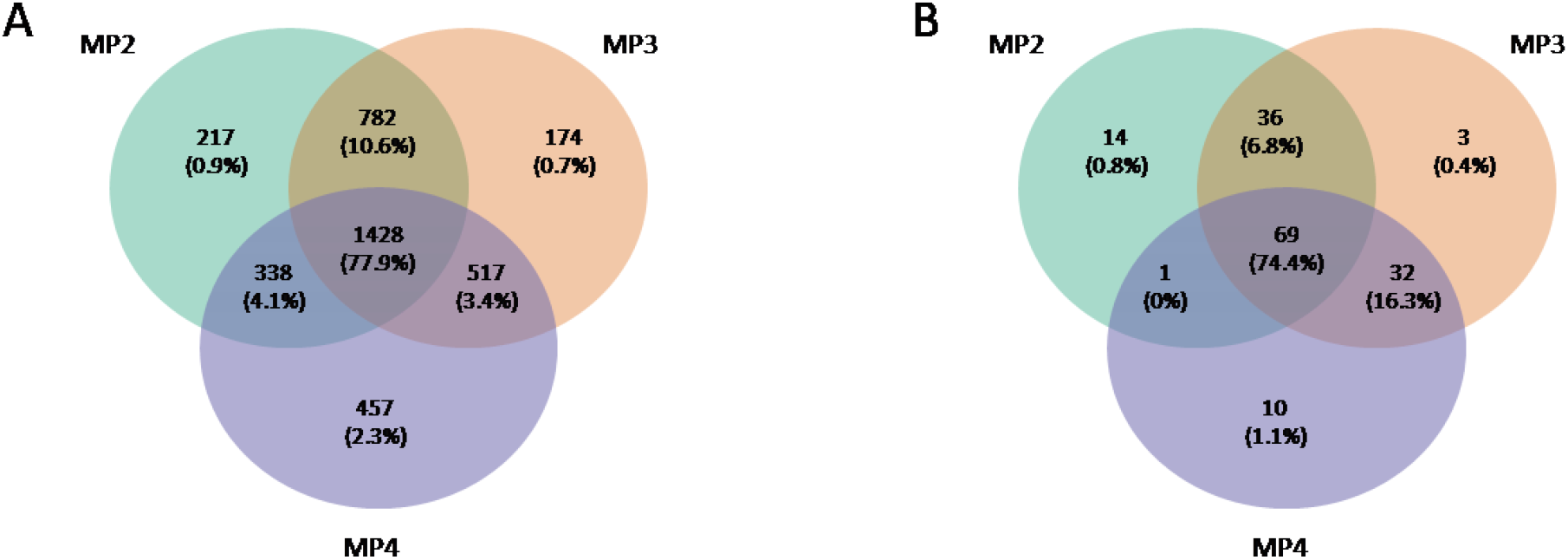
Venn diagram of the bacterial (A) and T4-type phages (B) ASVs. The numbers correspond to ASVs shared between two or three sites and the percentage indicated the proportion of sequences they account for.

**Supplementary Fig. 4.**
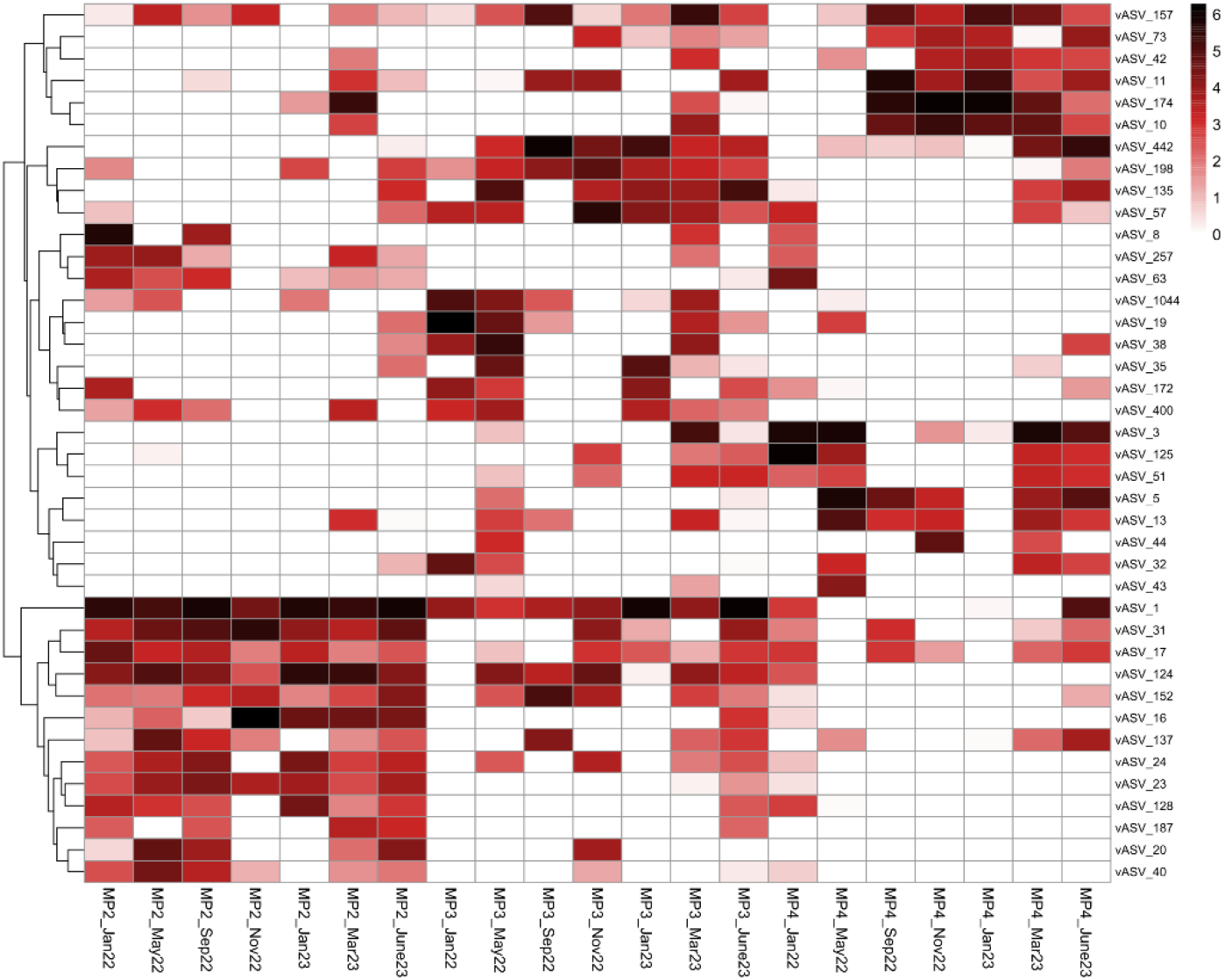
Hierarchical clustering using Euclidean distance and heatmap representation of relative abundance the 40 main viral ASVs after normalization by log-transformation. Site and sampling dates are shown on the horizontal axis and the vertical axis represents the normalized values of vASVs.

---

**Supplementary Table 1.** Numbers and sequence relative abundances of the shared T4-type phage ASVs (vASVs) between the three sites at the different sampling dates.

|  | January 22 |  | May 22 |  | September 22 |  | November 22 |  | January 23 |  | March 23 |  | June 23 |  |
| --- | --- | --- | --- | --- | --- | --- | --- | --- | --- | --- | --- | --- | --- | --- |
|  | Number of vASVs | relative abundance of viral sequences | Number of vASVs | relative abundance of viral sequences | Number of vASVs | relative abundance of viral sequences | Number of vASVs | relative abundance of viral sequences | Number of vASVs | relative abundance of viral sequences | Number of vASVs | relative abundance of viral sequences | Number of vASVs | relative abundance of viral sequences |
| MP2 | 21 | 2.5% | 31 | 13.9% | 40 | 31.5% | 16 | 5.9% | 14 | 2.7% | 32 | 9.6% | 17 | 3.4% |
| MP3 | 8 | 16.7% | 27 | 12% | 12 | 1.5% | 15 | 11.1% | 19 | 14.9% | 13 | 2.5% | 14 | 1.1% |
| MP4 | 13 | 11.6% | 13 | 7.5% | 10 | 7.6% | 16 | 60.8% | 15 | 24.3% | 19 | 4.2% | 11 | 1.2% |
| MP2&MP3 | 5 | 5.1% | 15 | 12.3% | 12 | 36.4% | 7 | 7.6% | 8 | 3.9% | 18 | 32.4% | 29 | 17.3% |
| MP2&MP4 | 22 | 43.2% | 3 | 5.1% | 3 | 12.3% | 1 | 0.1% | 1 | 24.3% | 6 | 3.2% | 6 | 1% |
| MP4&MP3 | 1 | 0.1% | 12 | 46.6% | 3 | 4.3% | 4 | 12.1% | 5 | 18.1% | 23 | 21.8% | 21 | 12.3% |
| MP2&MP4&MP3 | 3 | 20.6% | 3 | 2.6% | 2 | 6.5% | 2 | 2.5% | 1 | 11.7% | 10 | 26.4% | 23 | 63.6% |

## References

Anisimova, M., Gascuel, O., 2006. Approximate likelihood-ratio test for branches: A fast, accurate, and powerful alternative. Systematic Biology 55, 539–552. doi:10.1080/10635150600755453

Bastian, M., Heymann, S., Jacomy, M., 2009. Gephi: An Open Source Software for Exploring and Manipulating Networks. Proceedings of the Third International ICWSM Conference 361–362.

Bellas, C.M., Anesio, A.M., 2013. High diversity and potential origins of T4-type bacteriophages on the surface of Arctic glaciers. Extremophiles: Life under Extreme Conditions 17, 861–870. doi:10.1007/S00792-013-0569-X

Bethencourt, L., Bochet, O., Farasin, J., Aquilina, L., Borgne, T. Le, Quaiser, A., Biget, M., Michon-Coudouel, S., Labasque, T., Dufresne, A., 2020. Genome reconstruction reveals distinct assemblages of Gallionellaceae in surface and subsurface redox transition zones. FEMS Microbiology Ecology 96, 36. doi:10.1093/FEMSEC/FIAA036

Bi, L., Yu, D.T., Du, S., Zhang, L.M., Zhang, L.Y., Wu, C.F., Xiong, C., Han, L.L., He, J.Z., 2021. Diversity and potential biogeochemical impacts of viruses in bulk and rhizosphere soils. Environmental Microbiology 23, 588–599. doi:10.1111/1462-2920.15010

Braga, L.P.P., Spor, A., Kot, W., Breuil, M.-C., Hansen, L.H., Setubal, J.C., Philippot, L., 2020. Impact of phages on soil bacterial communities and nitrogen availability under different assembly scenarios. Microbiome 8, 52. doi:10.1186/s40168-020-00822-z

Breitbart, M., Rohwer, F., 2005. Here a virus, there a virus, everywhere the same virus? Trends in Microbiology 13, 278–284. doi:10.1016/J.TIM.2005.04.003/ASSET/96BE0BBA-5577-4565-83D4-E22930F7B80D/MAIN.ASSETS/GR3.JPG

Buchfink, B., Xie, C., Huson, D.H., 2014. Fast and sensitive protein alignment using DIAMOND. Nature Methods 2014 12:1 12, 59–60. doi:10.1038/NMETH.3176

Cahyani, V.R., Murase, J., Ishibashi, E., Asakawa, S., Kimura, M., 2009. T4-type bacteriophage communities estimated from the major capsid genes (g23) in manganese nodules in Japanese paddy fields. Soil Science and Plant Nutrition 55, 264–270. doi:10.1111/J.1747-0765.2009.00363.X

Cai, L., Xu, B., Li, H., Xu, Y., Wei, W., Zhang, R., 2023. Spatiotemporal Shift of T4-Like Phage Community Structure in the Three Largest Estuaries of China. Microbiology Spectrum 11. doi:10.1128/SPECTRUM.05203-22/SUPPL_FILE/SPECTRUM.05203-22-S0001.PDF

Callahan, B.J., McMurdie, P.J., Rosen, M.J., Han, A.W., Johnson, A.J.A., Holmes, S.P., 2016. DADA2: High-resolution sample inference from Illumina amplicon data. Nature Methods 2016 13:7 13, 581–583.

Carlson, H.K., Piya, D., Moore, M.L., Magar, R.T., Elisabeth, N.H., Deutschbauer, A.M., Arkin, A.P., Mutalik, V.K., 2023. Geochemical constraints on bacteriophage infectivity in terrestrial environments. ISME Communications 3. doi:10.1038/S43705-023-00297-7

Castledine, M., Buckling, A., 2024. Critically evaluating the relative importance of phage in shaping microbial community composition. Trends in Microbiology 957–969.

Chow, C.-E.E.T., Kim, D.Y., Sachdeva, R., Caron, D.A., Fuhrman, J.A., 2014. Top-down controls on bacterial community structure: microbial network analysis of bacteria, T4-like viruses and protists. The ISME Journal 8, 816–829. doi:10.1038/ismej.2013.199

Dar, N., Thompson, C.P., Williamson, K., 2023. Marker gene analysis reveals novel viral genetic diversity in unsaturated soils. Biology and Fertility of Soils 59, 139–151. doi:10.1007/s00374-022-01687-0

de Jonge, P.A., Nobrega, F.L., Brouns, S.J.J., Dutilh, B.E., 2019. Molecular and Evolutionary Determinants of Bacteriophage Host Range. Trends in Microbiology 27, 51–63. doi:10.1016/J.TIM.2018.08.006

Dedysh, S.N., Ivanova, A.A., 2019. Planctomycetes in boreal and subarctic wetlands: diversity patterns and potential ecological functions. FEMS Microbiology Ecology 95, 227. doi:10.1093/FEMSEC/FIY227

Dereeper, A., Guignon, V., Blanc, G., Audic, S., Buffet, S., Chevenet, F., Dufayard, J.F., Guindon, S., Lefort, V., Lescot, M., Claverie, J.M., Gascuel, O., 2008. Phylogeny.fr: robust phylogenetic analysis for the non-specialist. Nucleic Acids Research 36, W465. doi:10.1093/NAR/GKN180

Desplats, C., Krisch, H.M., 2003. The diversity and evolution of the T4-type bacteriophages. Research in Microbiology 154, 259–267. doi:10.1016/S0923-2508(03)00069-X

Emerson, J.B., Roux, S., Brum, J.R., Bolduc, B., Woodcroft, B.J., Jang, H. Bin, Singleton, C.M., Solden, L.M., Naas, A.E., Boyd, J.A., Hodgkins, S.B., Wilson, R.M., Trubl, G., Li, C., Frolking, S., Pope, P.B., Wrighton, K.C., Crill, P.M., Chanton, J.P., Saleska, S.R., Tyson, G.W., Rich, V.I., Sullivan, M.B., 2018. Host-linked soil viral ecology along a permafrost thaw gradient. Nature Microbiology 3, 870–880. doi:10.1038/s41564-018-0190-y

Erktan, A., Or, D., Scheu, S., 2020. The physical structure of soil: Determinant and consequence of trophic interactions. Soil Biology and Biochemistry 148, 107876. doi:10.1016/J.SOILBIO.2020.107876

Filée, J., Tétart, F., Suttle, C.A., Krisch, H.M., 2005. Marine T4-type bacteriophages, a ubiquitous component of the dark matter of the biosphere. Proceedings of the National Academy of Sciences of the United States of America 102, 12471–12476. doi:10.1073/PNAS.0503404102/SUPPL_FILE/03404FIG4.JPG

Foster, Z.S.L., Sharpton, T.J., Grünwald, N.J., 2017. Metacoder: An R package for visualization and manipulation of community taxonomic diversity data. PLOS Computational Biology 13, e1005404. doi:10.1371/JOURNAL.PCBI.1005404

Friedman, J., Alm, E.J., 2012. Inferring Correlation Networks from Genomic Survey Data. PLOS Computational Biology 8, e1002687. doi:10.1371/JOURNAL.PCBI.1002687

Fu, L., Niu, B., Zhu, Z., Wu, S., Li, W., 2012. CD-HIT: accelerated for clustering the next-generation sequencing data. Bioinformatics 28, 3150–3152. doi:10.1093/BIOINFORMATICS/BTS565

Fujihara, S., Murase, J., Tun, C.C., Matsuyama, T., Ikenaga, M., Asakawa, S., Kimura, M., 2010. Low diversity of T4-type bacteriophages in applied rice straw, plant residues and rice roots in Japanese rice soils: Estimation from major capsid gene (g23) composition. Soil Science &Plant Nutrition 56, 800–812. doi:10.1111/J.1747-0765.2010.00513.X

Fujii, T., Nakayama, N., Nishida, M., Sekiya, H., Kato, N., Asakawa, S., Kimura, M., 2008. Novel capsid genes (g23) of T4-type bacteriophages in a Japanese paddy field. Soil Biology and Biochemistry 40, 1049–1058. doi:10.1016/j.soilbio.2007.11.025

Gaillardet, J., Braud, I., Hankard, F., Anquetin, S., Bour, O., Dorfliger, N., de Dreuzy, J.R., Galle, S., Galy, C., Gogo, S., Gourcy, L., Habets, F., Laggoun, F., Longuevergne, L., Le Borgne, T., Naaim-Bouvet, F., Nord, G., Simonneaux, V., Six, D., Tallec, T., Valentin, C., Abril, G., Allemand, P., Arènes, A., Arfib, B., Arnaud, L., Arnaud, N., Arnaud, P., Audry, S., Comte, V.B., Batiot, C., Battais, A., Bellot, H., Bernard, E., Bertrand, C., Bessière, H., Binet, S., Bodin, J., Bodin, X., Boithias, L., Bouchez, J., Boudevillain, B., Moussa, I.B., Branger, F., Braun, J.J., Brunet, P., Caceres, B., Calmels, D., Cappelaere, B., Celle-Jeanton, H., Chabaux, F., Chalikakis, K., Champollion, C., Copard, Y., Cotel, C., Davy, P., Deline, P., Delrieu, G., Demarty, J., Dessert, C., Dumont, M., Emblanch, C., Ezzahar, J., Estèves, M., Favier, V., Faucheux, M., Filizola, N., Flammarion, P., Floury, P., Fovet, O., Fournier, M., Francez, A.J., Gandois, L., Gascuel, C., Gayer, E., Genthon, C., Gérard, M.F., Gilbert, D., Gouttevin, I., Grippa, M., Gruau, G., Jardani, A., Jeanneau, L., Join, J.L., Jourde, H., Karbou, F., Labat, D., Lagadeuc, Y., Lajeunesse, E., Lastennet, R., Lavado, W., Lawin, E., Lebel, T., Le Bouteiller, C., Legout, C., Lejeune, Y., Le Meur, E., Le Moigne, N., Lions, J., Lucas, A., Malet, J.P., Marais-Sicre, C., Maréchal, J.C., Marlin, C., Martin, P., Martins, J., Martinez, J.M., Massei, N., Mauclerc, A., Mazzilli, N., Molénat, J., Moreira-Turcq, P., Mougin, E., Morin, S., Ngoupayou, J.N., Panthou, G., Peugeot, C., Picard, G., Pierret, M.C., Porel, G., Probst, A., Probst, J.L., Rabatel, A., Raclot, D., Ravanel, L., Rejiba, F., René, P., Ribolzi, O., Riotte, J., Rivière, A., Robain, H., Ruiz, L., Sanchez-Perez, J.M., Santini, W., Sauvage, S., Schoeneich, P., Seidel, J.L., Sekhar, M., Sengtaheuanghoung, O., Silvera, N., Steinmann, M., Soruco, A., Tallec, G., Thibert, E., Lao, D.V., Vincent, C., Viville, D., Wagnon, P., Zitouna, R., 2018. OZCAR: the French network of critical zone observatories. Vadose Zone Journal 17, 0. doi:10.2136/vzj2018.04.0067

Goberna, M., Verdú, M., 2022. Cautionary notes on the use of co-occurrence networks in soil ecology. Soil Biology and Biochemistry 166, 108534. doi:10.1016/J.SOILBIO.2021.108534

Griffiths, R.I., Whiteley, A.S., O’Donnell, A.G., Bailey, M.J., 2000. Rapid method for coextraction of DNA and RNA from natural environments for analysis of ribosomal DNA- and rRNA-based microbial community composition. Applied and Environmental Microbiology 66, 5488–5491. doi:10.1128/aem.66.12.5488-5491.2000

Guindon, S., Dufayard, J.F., Lefort, V., Anisimova, M., Hordijk, W., Gascuel, O., 2010. New Algorithms and Methods to Estimate Maximum-Likelihood Phylogenies: Assessing the Performance of PhyML 3.0. Systematic Biology 59, 307–321. doi:10.1093/SYSBIO/SYQ010

Guseva, K., Darcy, S., Simon, E., Alteio, L. V., Montesinos-Navarro, A., Kaiser, C., 2022. From diversity to complexity: Microbial networks in soils. Soil Biology and Biochemistry 169, 108604. doi:10.1016/J.SOILBIO.2022.108604

He, M., Cai, L., Zhang, C., Jiao, N., Zhang, R., 2017. Phylogenetic diversity of T4-type phages in sediments from the subtropical Pearl River Estuary. Frontiers in Microbiology 8, 260221. doi:10.3389/FMICB.2017.00897/BIBTEX

Hedrich, S., Schlömann, M., Johnson, D.B., 2011. The iron-oxidizing proteobacteria. Microbiology (Reading, England) 157, 1551–64. doi:10.1099/mic.0.045344-0

Hendrix, R.W., Smith, M.C.M., Burns, R.N., Ford, M.E., Hatfull, G.F., 1999. Evolutionary relationships among diverse bacteriophages and prophages: All the world’s a phage. Proceedings of the National Academy of Sciences of the United States of America 96, 2192–2197. doi:10.1073/PNAS.96.5.2192/ASSET/A9620A61-08D5-4067-A7BC-8FA499701923/ASSETS/GRAPHIC/PQ0594730004.JPEG

Hu, H.W., Chen, D., He, J.Z., 2015. Microbial regulation of terrestrial nitrous oxide formation: understanding the biological pathways for prediction of emission rates. FEMS Microbiology Reviews 39, 729–749. doi:10.1093/FEMSRE/FUV021

Ivanova, A.A., Oshkin, I.Y., Danilova, O. V., Philippov, D.A., Ravin, N. V., Dedysh, S.N., 2022. Rokubacteria in northern peatlands: Habitat preferences and diversity patterns. Microorganisms 10. doi:10.3390/MICROORGANISMS10010011

Jamindar, S., Polson, S.W., Srinivasiah, S., Waidner, L., Wommack, K.E., 2012. Evaluation of two approaches for assessing the genetic similarity of virioplankton populations as defined by genome size. Applied and Environmental Microbiology 78, 8773–8783. doi:10.1128/AEM.02432-12/SUPPL_FILE/ZAM999103941SO2.PDF

Jansson, J.K., Wu, R., 2022. Soil viral diversity, ecology and climate change. Nature Reviews Microbiology 2022 21:5 21, 296–311. doi:10.1038/S41579-022-00811-Z

Kang, W., Xiao, Y., Li, W., Cheng, A., Cheng, C., Jia, Z., Yu, L., 2024. Paddy cultivation in degraded karst wetland soil can significantly improve the physiological and ecological functions of carbon-fixing resident microorganisms. Science of The Total Environment 909, 168187. doi:10.1016/J.SCITOTENV.2023.168187

Klindworth, A., Pruesse, E., Schweer, T., Peplies, J., Quast, C., Horn, M., Glöckner, F.O., 2013. Evaluation of general 16S ribosomal RNA gene PCR primers for classical and next-generation sequencing-based diversity studies. Nucleic Acids Research 41, 1–11. doi:10.1093/nar/gks808

Krisch, H.M., Comeau, A.M., 2008. The immense journey of bacteriophage T4—From d’Hérelle to Delbrück and then to Darwin and beyond. Research in Microbiology 159, 314–324. doi:10.1016/J.RESMIC.2008.04.014

Kuzyakov, Y., Mason-Jones, K., 2018. Viruses in soil: Nano-scale undead drivers of microbial life, biogeochemical turnover and ecosystem functions. Soil Biology and Biochemistry 127, 305–317. doi:10.1016/J.SOILBIO.2018.09.032

Lan, J., Wang, S., Wang, J., Qi, X., Long, Q., Huang, M., 2022. The Shift of Soil Bacterial Community After Afforestation Influence Soil Organic Carbon and Aggregate Stability in Karst Region. Frontiers in Microbiology 13, 901126. doi:10.3389/FMICB.2022.901126/BIBTEX

Lenferink, W.B., van Alen, T.A., Jetten, M.S.M., Op den Camp, H.J.M., van Kessel, M.A.H.J., Lücker, S., 2024. Genomic analysis of the class Phycisphaerae reveals a versatile group of complex carbon-degrading bacteria. Antonie Van Leeuwenhoek 117, 104. doi:10.1007/S10482-024-02002-7

Letunic, I., Bork, P., 2024. Interactive Tree of Life (iTOL) v6: recent updates to the phylogenetic tree display and annotation tool. Nucleic Acids Research 52, W78–W82. doi:10.1093/NAR/GKAE268

Li, H., Liu, L., Wang, Y., Cai, L., He, M., Wang, L., Hu, C., Jiao, N., Zhang, R., 2021. T4-like myovirus community shaped by dispersal and deterministic processes in the South China Sea. Environmental Microbiology 23, 1038–1052. doi:10.1111/1462-2920.15290

Li, Yong, Liu, H., Pan, H., Zhu, X., Liu, C., Zhang, Q., Luo, Y., Di, H., Xu, J., 2019. T4-type viruses: Important impacts on shaping bacterial community along a chronosequence of 2000-year old paddy soils. Soil Biology and Biochemistry 128, 89–99. doi:10.1016/j.soilbio.2018.10.007

Li, Yaying, Pan, F., Yao, H., 2019. Response of symbiotic and asymbiotic nitrogen-fixing microorganisms to nitrogen fertilizer application. Journal of Soils and Sediments 19, 1948–1958. doi:10.1007/S11368-018-2192-Z/METRICS

Liu, C., Cui, Y., Li, X., Yao, M., 2021. microeco: an R package for data mining in microbial community ecology. FEMS Microbiology Ecology 97, 255. doi:10.1093/FEMSEC/FIAA255

Liu, J., Wang, G., Zheng, C., Yuan, X., Jin, J., Liu, X., 2011. Specific assemblages of major capsid genes (g23) of T4-type bacteriophages isolated from upland black soils in Northeast China. Soil Biology and Biochemistry 43, 1980–1984. doi:10.1016/J.SOILBIO.2011.05.005

Liu, J., Yu, Z., Wang, X., Jin, J., Liu, X., Wang, G., 2016. The distribution characteristics of the major capsid gene (g23) of T4-type phages in paddy floodwater in Northeast China. Soil Science and Plant Nutrition 62, 133–139. doi:10.1080/00380768.2016.1163507

Liu, Junjie, Wang, G., Wang, Q., Liu, Judong, Jin, J., Liu, X., 2012. Phylogenetic diversity and assemblage of major capsid genes (g23) of T4-type bacteriophages in paddy field soils during rice growth season in Northeast China. Soil Science and Plant Nutrition 58, 435–444. doi:10.1080/00380768.2012.703610

Liu, L., Cai, L., Zhang, R., 2017. Co-existence of freshwater and marine T4-like myoviruses in a typical subtropical estuary. FEMS Microbiology Ecology 93, 119. doi:10.1093/FEMSEC/FIX119

López-Bueno, A., Tamames, J., Velázquez, D., Moya, A., Quesada, A., Alcamí, A., 2009. High diversity of the viral community from an Antarctic lake. Science 326, 858–861. doi:10.1126/SCIENCE.1179287/SUPPL_FILE/LOPEZ-BUENO.SOM.PDF

Lyu, Z., Shao, N., Akinyemi, T., Whitman, W.B., 2018. Methanogenesis. Current Biology: CB 28, R727–R732. doi:10.1016/J.CUB.2018.05.021

Ma, B., Wang, Y., Zhao, K., Stirling, E., Lv, X., Yu, Y., Hu, L., Tang, C., Wu, C., Dong, B., Xue, R., Dahlgren, R.A., Tan, X., Dai, H., Zhu, Y.G., Chu, H., Xu, J., 2024. Biogeographic patterns and drivers of soil viromes. Nature Ecology & Evolution 2024 8:4 8, 717–728. doi:10.1038/S41559-024-02347-2

Martin, M., 2011. Cutadapt removes adapter sequences from high-throughput sequencing reads. EMBnet.Journal 17.

McMurdie, P.J., Holmes, S., 2013. Phyloseq: An R Package for Reproducible Interactive Analysis and Graphics of Microbiome Census Data. PLoS ONE 8. doi:10.1371/journal.pone.0061217

Monard, C., Gantner, S., Bertilsson, S., Hallin, S., Stenlid, J., 2016. Habitat generalists and specialists in microbial communities across a terrestrial-freshwater gradient. Scientific Reports 6, 37719. doi:10.1038/srep37719

Nakayama, N., Asakawa, S., Kimura, M., 2009. Comparison of g23 gene sequence diversity between Novosphingobium and Sphingomonas phages and phage communities in the floodwater of a Japanese paddy field. Soil Biology and Biochemistry 41, 179–185. doi:10.1016/J.SOILBIO.2008.06.008

Nicolaisen, M.H., Baelum, J., Jacobsen, C.S., Sorensen, J., 2008. Transcription dynamics of the functional tfdA gene during MCPA herbicide degradation by Cupriavidus necator AEO106 (pRO101) in agricultural soil. Environmental Microbiology 10, 571–579. doi:doi:10.1111/j.1462-2920.2007.01476.x

Nicolas, A.M., Sieradzki, E.T., Pett-Ridge, J., Banfield, J.F., Taga, M.E., Firestone, M.K., Blazewicz, S.J., 2022. Isotope-enrichment reveals active viruses follow microbial host dynamics during rewetting of a California grassland soil. BioRxiv 2022.09.30.510406. doi:10.1101/2022.09.30.510406

Nunes, F.L.D., Aquilina, L., De Ridder, J., Francez, A.J., Quaiser, A., Caudal, J.P., Vandenkoornhuyse, P., Dufresne, A., 2015. Time-scales of hydrological forcing on the geochemistry and bacterial community structure of temperate peat soils. Scientific Reports 2015 5:1 5, 1–11. doi:10.1038/SREP14612

Oksanen, J., Blanchet, F.G., Kindt, R., Legendre, P., Minchin, P.R., O’Hara, R.B., Simpson, G.L., Solymos, P., Stevens, M.H.H., Wagner, H., 2015. vegan: Community Ecology Package.

Olson, S.A., 2002. Emboss opens up sequence analysis. Briefings in Bioinformatics 3, 87–91. doi:10.1093/bib/3.1.87

Peres, F. V., Paula, F.S., Bendia, A.G., Gontijo, J.B., de Mahiques, M.M., Pellizari, V.H., 2023. Assessment of prokaryotic communities in Southwestern Atlantic deep-sea sediments reveals prevalent methanol-oxidising Methylomirabilales. Scientific Reports 2023 13:1 13, 1–12. doi:10.1038/S41598-023-39415-9

Petrov, V.M., Ratnayaka, S., Nolan, J.M., Miller, E.S., Karam, J.D., 2010. Genomes of the T4-related bacteriophages as windows on microbial genome evolution. Virology Journal 7, 1–19. doi:10.1186/1743-422X-7-292/TABLES/5

Poulter, B., Fluet-Chouinard, E., Hugelius, G., Koven, C., Fatoyinbo, L., Page, S.E., Rosentreter, J.A., Smart, L.S., Taillie, P.J., Thomas, N., Zhang, Z., Wijedasa, L.S., 2021. A review of global wetland carbon stocks and management challenges. Wetland Carbon and Environmental Management 1–20. doi:10.1002/9781119639305.CH1

Pratama, A.A., van Elsas, J.D., 2018. The neglected’ soil virome – potential role and impact. Trends in Microbiology 26, 649–662. doi:10.1016/j.tim.2017.12.004

Rice, P., Longden, L., Bleasby, A., 2000. EMBOSS: The European Molecular Biology Open Software Suite. Trends in Genetics 16, 276–277. doi:10.1016/S0168-9525(00)02024-2

Roux, S., Enault, F., Ravet, V., Pereira, O., Sullivan, M.B., 2015. Genomic characteristics and environmental distributions of the uncultivated Far-T4 phages. Frontiers in Microbiology 6, 128504. doi:10.3389/FMICB.2015.00199/BIBTEX

Starr, E.P., Nuccio, E.E., Pett-Ridge, J., Banfield, J.F., Firestone, M.K., 2019. Metatranscriptomic reconstruction reveals RNA viruses with the potential to shape carbon cycling in soil. Proceedings of the National Academy of Sciences of the United States of America 116, 25900–25908. doi:10.1073/PNAS.1908291116/SUPPL_FILE/PNAS.1908291116.SAPP.PDF

Subedi, D., Barr, J.J., 2021. Temporal Stability and Genetic Diversity of 48-Year-Old T-Series Phages. MSystems 6. doi:10.1128/MSYSTEMS.00990-20

Sullivan, M.B., Huang, K.H., Ignacio-Espinoza, J.C., Berlin, A.M., Kelly, L., Weigele, P.R., DeFrancesco, A.S., Kern, S.E., Thompson, L.R., Young, S., Yandava, C., Fu, R., Krastins, B., Chase, M., Sarracino, D., Osburne, M.S., Henn, M.R., Chisholm, S.W., 2010. Genomic analysis of oceanic cyanobacterial myoviruses compared with T4-like myoviruses from diverse hosts and environments. Environmental Microbiology 12, 3035–3056. doi:10.1111/J.1462-2920.2010.02280.X

Suttle, C.A., 2005. Viruses in the sea. Nature 437, 356–361. doi:10.1038/nature04160

Tang, H., Xiao, X., Li, C., Tang, W., Cheng, K., Wang, K., Pan, X., Li, W., 2019. Effects of Rhizosphere and Long-Term Fertilization Practices on the Activity and Community Structure of Denitrifiers Under Double-Cropping Rice Field. Communications in Soil Science and Plant Analysis 50, 682–697. doi:10.1080/00103624.2019.1589480

Tetart, F., Desplats, C., Kutateladze, M., Monod, C., Ackermann, H.W., Krisch, H.M., 2001. Phylogeny of the Major Head and Tail Genes of the Wide-Ranging T4-Type Bacteriophages. Journal of Bacteriology 183, 358. doi:10.1128/JB.183.1.358-366.2001

Unger, I.M., Kennedy, A.C., Muzika, R.M., 2009. Flooding effects on soil microbial communities. Applied Soil Ecology 42, 1–8. doi:10.1016/J.APSOIL.2009.01.007

Wang, G., Hayashi, M., Saito, M., Tsuchiya, K., Asakawa, S., Kimura, M., 2009a. Survey of major capsid genes (g23) of T4-type bacteriophages in Japanese paddy field soils. Soil Biology and Biochemistry 41, 13–20. doi:10.1016/J.SOILBIO.2008.07.008

Wang, G., Jin, J., Asakawa, S., Kimura, M., 2009b. Survey of major capsid genes (g23) of T4-type bacteriophages in rice fields in Northeast China. Soil Biology and Biochemistry 41, 423–427. doi:10.1016/J.SOILBIO.2008.11.012

Wang, G., Yu, Z., Liu, J., Jin, J., Liu, X., Kimura, M., 2011. Molecular analysis of the major capsid genes (g23) of T4-type bacteriophages in an upland black soil in Northeast China. Biology and Fertility of Soils 47, 273–282. doi:10.1007/S00374-010-0533-1/FIGURES/4

Wang, P., Ouyang, W., Zhu, W., Geng, F., Tulcan, R.X.S., Lin, C., 2023. Wetland soil carbon dioxide emission dynamics with external dissolved organic matter in mid–high-latitude forested watershed. Agricultural and Forest Meteorology 333, 109381. doi:10.1016/J.AGRFORMET.2023.109381

Wang, X., Tang, Y., Yue, X., Wang, S., Yang, K., Xu, Y., Shen, Q., Friman, V.P., Wei, Z., 2024. The role of rhizosphere phages in soil health. FEMS Microbiology Ecology 100. doi:10.1093/FEMSEC/FIAE052

Weinbauer, M.G., 2004. Ecology of prokaryotic viruses. FEMS Microbiology Reviews 28, 127–181. doi:10.1016/j.femsre.2003.08.001

Wickham, H., 2016. ggplot2: Elegant graphics for data analysis, Use R! Springer Verlag. doi:10.1007/978-3-319-24277-4

Wilhelm, S.W., Suttle, C.A., 1999. Viruses and nutrient cycles in the sea. BioScience 49, 781–788. doi:10.2307/1313569

Williamson, K.E., Fuhrmann, J.J., Wommack, K.E., Radosevich, M., 2017. Viruses in soil ecosystems: An unknown quantity within an unexplored territory. Annual Review of Virology 4, 201–219. doi:10.1146/annurev-virology-101416-041639

Wu, X., Peng, J., Liu, P., Bei, Q., Rensing, C., Li, Y., Yuan, H., Liesack, W., Zhang, F., Cui, Z., 2021. Metagenomic insights into nitrogen and phosphorus cycling at the soil aggregate scale driven by organic material amendments. Science of The Total Environment 785, 147329. doi:10.1016/J.SCITOTENV.2021.147329

Yilmaz, P., Parfrey, L.W., Yarza, P., Gerken, J., Pruesse, E., Quast, C., Schweer, T., Peplies, J., Ludwig, W., Glöckner, F.O., 2014. The SILVA and “All-species Living Tree Project (LTP)” taxonomic frameworks. Nucleic Acids Research 42. doi:10.1093/NAR/GKT1209

Zhang, J., Liu, S., Liu, C., Wang, H., Luan, J., Liu, X., Guo, X., Niu, B., 2021. Different mechanisms underlying divergent responses of autotrophic and heterotrophic respiration to long-term throughfall reduction in a warm-temperate oak forest. Forest Ecosystems 2021 8:1 8, 1–11. doi:10.1186/S40663-021-00321-Z

Zheng, C., Wang, G., Liu, J., Song, C., Gao, H., Liu, X., 2013. Characterization of the Major Capsid Genes (g23) of T4-Type Bacteriophages in the Wetlands of Northeast China. Microbial Ecology 65, 616–625. doi:10.1007/s00248-012-0158-z

Zhou, Z., Liang, X., Zhang, N., Xie, N., Huang, Y., Zhou, Y., Li, B., 2024. The impact of soil viruses on organic carbon mineralization and microbial biomass turnover. Applied Soil Ecology 202, 105554. doi:10.1016/J.APSOIL.2024.105554

